# *Prpf8^N1531S^* homozygous mutant mouse embryos have multiple defects in cardiac development and show aberrant splicing of the cardiac transcription factor *Tead1*

**DOI:** 10.64898/2026.04.17.719138

**Authors:** Wasay Mohiuddin Shaikh Qureshi, Haoxiang Zhou, Abigail Bennington, Nouf Althali, Anna van der Zwaluw, Michael Boylan, Louise A Stephen, Kashish Jain, Basudha Basu, Dapeng Wang, Colin A Johnson, Kathryn E Hentges

**Author notes:** **Corresponding author:** Kathryn E Hentges. MB: Developmental Biology and Cancer Department, UCL Great Ormond Street Institute of Child Health, London WC1N 1EH, UK LAS: Division of Functional Genetics, The Roslin Institute, The University of Edinburgh, Midlothian, EH25 9RG UK. KJ: Department of Molecular Biology and Genetics, Cornell University, Ithaca, New York, 14850, USA DW: Shandong Key Laboratory of Intelligent Oil & Gas Industrial Software, Qingdao Institute of Software, College of Computer Science and Technology, China University of Petroleum (East China), Qingdao, 266580, China.

## Abstract

Mutations in the spliceosomal gene *PRPF8* are associated with a range of human diseases. Studies in mouse and zebrafish suggest that *Prpf8* also has a developmental function. Here, using a *Prpf8* mutant mouse line isolated from a chemical induced mutagenesis screen, we uncover a previously unrecognised and essential role for *Prpf8* in heart development, consistent with the embryonic lethality observed in *Prpf8^N1531S^* homozygous mutants. *Prpf8^N1531S^* mutant embryos display severe defects in ventricular trabeculation and compact zone formation, accompanied by increased cardiomyocyte proliferation specifically in the compact zone. Mutant embryonic hearts also exhibit disrupted cellular organisation, altered cytoskeletal architecture and changes in extracellular matrix protein expression. Notably, these cardiac abnormalities were exacerbated in embryos exhibiting cardiac looping defects. Transcriptomic analysis identified multiple aberrantly spliced transcripts in *Prpf8^N1531S^* mutant embryos, among which the cardiac transcription factor Tead1 was selected as a key functional candidate due to it known role in cardiac ventricle wall developemnt. *Tead1* mis-splicing generated an in-frame, lower molecular weight protein isoform, associated with reduced overall TEAD1 expression. The *Tead1* mis-spliced isoform showed altered nuclear localisation and dysregulation of TEAD1-dependent gene network important for heart development, including known cardiac sarcomeric genes. In addition, we observed reduced levels of the intracellular domain of the NOTCH1 receptor (NICD1), indicating impaired Notch signalling.. These findings suggest that impaired TEAD1-dependent transcription and Notch signalling contribute to abnormal cardiac trabeculation and compact zone development, highlighting a critical role for *Prpf8* in maintaining proper heart development through the regulation of cardiac transcription factor expression and associated signalling networks. This study offers new mechanistic insights into congenital heart diseases linked to spliceosomal gene mutations.

## Introduction

Cardiac development is a complex and tightly regulated process, driven by the coordinated interplay of transcription factors, signalling pathways, and epigenetic regulators. Aberrations in these components can lead to congenital heart disease (CHD), the most prevalent type of birth defect, affecting approximately 1% of live births globally (Yue et al., 2021, Ma et al., 2024, Wu et al., 2020, van der Linde et al., 2011). Advances in genomic analysis have identified numerous genes involved in CHD pathogenesis (Chhatwal et al., 2023). However, 70–80% of CHD cases remain unexplained by known genetic or environmental factors (Cowan and Ware, 2015). Given the high prevalence and potentially life-threatening severity of CHD, identifying the genetic and molecular mechanisms underlying cardiac morphogenesis is essential for improving diagnosis, patient management, and genetic and genomic counselling.

Mutations in genes involved in fundamental cellular processes, including RNA splicing, have increasingly been implicated in CHD (Jang et al., 2023, Mehta and Touma, 2023, Engal et al., 2024, Richards and Garg, 2010, Spielmann et al., 2022). The spliceosomal machinery plays a vital role in regulating gene expression by generating transcript diversity and enabling tissue-specific isoform expression (Wilkinson et al., 2022, Manuel et al., 2023, García-Moreno and Romão, 2020). Mis-splicing can produce abnormal protein isoforms that lack essential functional domains or acquire deleterious properties (Jang et al., 2023, Mehta and Touma, 2023, Engal et al., 2024). During heart development, alternative splicing fine-tunes the expression of key cardiac transcription factors and signalling components to regulate cardiac morphogenesis. Disruption of splicing fidelity through mutations in spliceosomal components, such as PRPF8 (Jiang et al., 2025), RBM20 (Guo et al., 2012), RBFOX2 (Verma et al., 2022), EFTUD2 (Lines et al., 2012, Yang et al., 2022), SF3B4 (Bernier et al., 2012), TXNL4A (Goos et al., 2017) or SNRPB (Lynch et al., 2014) can lead to aberrant transcript isoform production, thereby perturbing developmental signalling pathways and contributing to CHD.

Through an ENU (N-ethyl-N-nitrosourea) mouse mutagenesis screen, a homozygous embryonic-lethal mutation in the spliceosomal gene *pre-mRNA processing factor 8* (*Prpf8*) was identified (Jiang et al., 2025). This mutation results in a missense substitution of asparagine with serine at amino acid 1531 (*Prpf8^N1531S^*). PRPF8 is ubiquitously expressed and has a role in pre-mRNA splicing as core component of precatalytic, catalytic and postcatalytic spliceosomal complexes, both of the predominant U2-type spliceosome and the minor U12-type spliceosome (Atkinson et al., 2024, Wahl et al., 2009). PRPF8 functions as a scaffold that mediates the ordered assembly of spliceosomal proteins and snRNAs, and is required for the assembly of the U4/U6-U5 tri-snRNP complex. This acts as a scaffold that positions spliceosomal U2, U5 and U6 snRNAs at splice sites on pre-mRNA substrates, so that splicing can occur (Galej et al., 2013, Grainger and Beggs, 2005, Nguyen et al., 2015). Consistent with its essential role, *Prpf8* null mouse embryos die during early development (Graziotto et al., 2011), making hypomorphic alleles such as *Prpf8^N1531S^* uniquely valuable for studying spliceosomal function *in vivo*. Pathogenic PRPF8 variants have been reported in autosomal dominant retinitis pigmentosa and neurodevelopmental disorders, indicating that partial impairment of PRPF8-mediated splicing can produce tissue-specific disease phenotypes (Růžičková and Staněk, 2017, O’Grady et al., 2022). Similarly, in zebrafish, the *cephalophonus* (*cph*) mutant harbouring a truncating mutation in *prpf8* exhibits global splicing defects with cardiac looping abnormalities (Keightley et al., 2013, Jiang et al., 2025). More recently, we have identified a role for PRPF8 in left–right patterning in mouse embryos, along with cardiac developmental defects (Jiang et al., 2025). However, despite these observations, the requirement for *Prpf8* in cardiac chamber development has not been examined.

Given the essential role of PRPF8 in maintaining splicing fidelity, we hypothesised that PRPF8 disruption may alter the splicing of key cardiac regulators. Transcriptomic analysis of *Prpf8^N1531S^* homozygous embryos (Jiang et al., 2025) identified multiple mis-splicing events, among which the cardiac transcription factor *Tead1* was selected as a key functional candidate due to its known role in ventricle wall development (Chen et al., 1994, Wen et al., 2019). We propose that aberrant splicing of *Tead1* in *Prpf8^N1531S^* homozygous embryos generates an in-frame protein isoform with reduced transcriptional activity, thereby limiting TEAD1-dependent gene regulation in mutant hearts. These findings support a model in which partial disruption of PRPF8-mediated splicing impairs TEAD1 function, leading to altered signalling networks and defective cardiac chamber development.

## Methods

### *Prpf8^N1531S^* mouse line, genotyping and embryo dissection

The *Prpf8^N1531S^* strain, which carries the chromosomal inversion Inv(11)8BrdTrp-Wnt3129s, was previously reported as the *l11Jus27* strain isolated from an ENU mutagenesis screen (Kile et al., 2003). We identified the causative mutation in this line as *Prpf8^N1531S^* (Jiang et al., 2025). These mice were maintained on a 129S5/SvEvBrd background. The mice were housed at University of Manchester, UK, in the AAALAC-accredited Biological Services Facility, in a controlled environment under a 12-hour light/dark cycle, at a temperature of 20–24°C and relative humidity of 45–65%, with food and water provided *ad libitum*. Animals were maintained in individually ventilated cages with appropriate environmental enrichment. All procedures were carried out in accordance with UK Home Office regulations (project licence number PP3720525) and local ethical approval from the Institutional Animal Welfare and Ethical Review Body. Efforts were made to minimise animal suffering, and all procedures were designed to reduce the number of animals used and their distress. Timed mating was confirmed by the presence of a vaginal plug, with embryonic day (E) 0.5 defined as noon on the day the vaginal plug was detected. Mice were euthanised using a Schedule 1 cervical dislocation method following UK Home Office regulations. Embryos were dissected from timed-mated pregnant female mice in ice-cold phosphate buffered saline (PBS), and their morphology was analysed using a Leica dissecting microscope. Genotyping was performed using primers targeting microsatellite regions to differentiate between C57BL/6 (mutant) and 129S5 (wild-type (WT)) alleles as described previously (Ridge et al., 2017) (Genotyping primers sequences, Forward GCGACTTATCTTCTACATGGGG and Reverse GGGGACGGAGGGCTTTATT). All WT embryos were derived from 129S5/SvEvBrd mouse line.

### Immunofluorescence staining and quantitative analysis

Mouse embryos or embryonic hearts were fixed overnight in 4% paraformaldehyde at 4°C. Fixed tissues were cryoprotected in 30% sucrose, embedded in OCT compound, and sectioned at a thickness of 10 µm. Antigen retrieval was performed on tissue sections using a solution containing 10 mM tri-sodium citrate and 0.05% Tween-20 (pH 6.0). This was followed by permeabilisation for 1 hour with 0.3% Triton X-100 in PBS (0.3% PBT) and blocking for 1 hour at room temperature using SuperBlock™ Blocking Buffer (Thermo Fisher Scientific, 37580).

Sections were incubated overnight at 4°C with primary antibodies (Supplementary Table 1) diluted in primary antibody diluent (Merck, KP31812). The following day, sections were washed with 0.1% PBT, and secondary antibodies conjugated to Alexa Fluor dyes or Phalloidin or Wheat Germ Agglutinin (WGA) stain (Supplementary Table 1), diluted in secondary antibody diluent (Merck, KP31855), were applied for 1 hour at room temperature, followed by additional washes in PBT. DAPI was used for nuclear staining. Slides were mounted using ProLong™ Gold Antifade Mountant (Thermo Fisher Scientific, P36934) and imaged using a Leica SP8 inverted confocal microscope.

Using ImageJ software, midline sections of E10.5 hearts (displaying all four cardiac chambers) stained with the cardiomyocyte marker MF20, PECAM and DAPI were analysed to measure trabeculation length, trabeculae number, and compact zone thickness in the left and right ventricles. Trabeculation length was defined as the linear distance from the base to the tip of each trabeculae. Compact zone thickness was measured as the distance between the epicardial surface and the base of the trabeculae. Ten measurements of compact zone thickness were taken per ventricle for each section analysed.

Proliferation of cardiac cells was assessed by immunofluorescence staining for the proliferation marker phospho-histone H3 (PH3). The number of PH3-positive cells was quantified across all cardiac cell types. The proliferation index was calculated as the percentage of PH3-positive cells relative to the total number of DAPI-stained nuclei.

### RT-PCR, TA Cloning and Sanger Sequencing

Total RNA was extracted from E10.5 embryos or embryonic hearts using TRIzol reagent according to the manufacturer’s protocol (Invitrogen; catalogue number 15596026). For complementary DNA (cDNA) synthesis, we used the High-Capacity cDNA Reverse Transcription Kit (Applied Biosystems) following the manufacturer’s instructions. RT-PCR was conducted using PCR Biosciences Ultra Mix Red polymerase with primers designed to amplify target *Tead1* exons (Supplementary Table 2). PCR products were separated by agarose gel electrophoresis and visualised under a UV transilluminator to identify the expected DNA bands. Amplified *Tead1* templates were inserted into the pCR^TM^2.1 vector provided in the TA cloning® Kit (Thermo Fisher Scientific). The ligation reaction was set up according to the manufacturer’s protocol, and the ligated products were transformed into One Shot® Mach1™-T1R chemically competent *E. coli* cells. For Sanger sequencing, the DNA band of interest was excised from the gel, and DNA extraction was performed using the Monarch® DNA Gel Extraction Kit (New England Biolabs). The extracted DNA samples or plasmids were diluted to the appropriate concentration and sequenced using Eurofins Genomics sequencing facility. The sequencing chromatograms were analysed using SnapGene software, and the resulting sequences were aligned against the mouse genome (GRCm39/mm39) to confirm the identity of the amplified sequences.

### Whole-mount *In Situ* Hybridisation

For digoxigenin (DIG)-labelled *in situ* hybridisation probe synthesis, we used either plasmid (Irx4) or PCR templates, with the gene-specific primer sequences listed in supplementary Table 3. Whole-mount *in situ* hybridisation was performed according to established protocols (Henrique et al., 1995). In brief, the mouse embryos were first fixed overnight in 4% paraformaldehyde at 4°C. The embryos were then hybridised with the DIG-labelled probes overnight at an optimised temperature. To detect the hybridised probes, the embryos were incubated with alkaline phosphatase-conjugated anti-DIG antibodies. The staining reaction was initiated using NBT/BCIP substrate solution, to develop the colourimetric signal. This process was carefully monitored, and the reaction was stopped once the desired staining intensity was achieved. Finally, the stained embryos were washed and imaged using a dissecting microscope.

### TEAD1 FLAG-tagged plasmid generation, HEK293 cell transfection and western blot analysis

Based on RNA sequencing data and Sanger sequencing results, three mouse *Tead1* FLAG-tagged CMV promoter plasmids (WT and two splice variants) were synthesised by VectorBuilder. The plasmids were transfected into HEK293T cells according to the method described previously (Wood et al., 2019). Briefly, HEK293T cells were seeded at a density of 0.25 × 10^6^ cells per well in 6-well plates. The following day, using FuGENE® HD transfection reagent (Promega), the cells were transfected with 2 µg of plasmids containing either the WT or splice variant sequence. At 24 h post-transfection, the cells were harvested for western blot analysis.

For western blot analysis, protein lysate were prepared from E10.5 mouse embryos or cultured HEK cells using RIPA buffer (Sigma-Aldrich R0278) and protease inhibitors cocktail (Promega G6521). The proteins were separated by SDS-PAGE and transferred to PVDF membranes as described (Wood et al., 2019). Membranes were probed with antibodies against TEAD1 (Cell Signaling Technology, (D9X2L) #12292; 1:1000), YAP1 (Proteintech, 13584-1-AP; 1:1000), NICD1 (Cell signalling 7194; 1:1000), FLAG (Sigma-Aldrich, F3165; 1:1000), and GAPDH (Proteintech, 10494-1-AP; 1:10,000; loading control), followed by incubation with fluorescence-conjugated LI-COR secondary antibodies (IRDye® 800CW goat anti-mouse IgG, 926-32210, and IRDye® 680RD goat anti-rabbit IgG, 926-68071). The fluorescence signals were detected using the Odyssey Infrared Imaging System (LI-COR Biosciences) and protein bands were analysed using Image Studio^TM^ software.

### H9c2 cells culture, transfection and immunostaining

The embryonic rat ventricular tissue derived cardiomyoblast cell line H9c2 was used to examine the effects of mis-spliced TEAD1 isoforms on nuclear localisation. H9c2 cells were seeded at 2.5 × 10⁴ cells per well in 24-well plates containing sterile glass coverslips (Watkins et al., 2011). After 24 h, cells were transfected with the TEAD1 plasmid using Lipofectamine 3000 (Thermo Fisher Scientific) according to the manufacturer’s instructions. At 48 h post-transfection, cells were washed with PBS, fixed, permeabilised, and blocked, followed by overnight incubation with the appropriate primary antibody. Cells were then washed and incubated with fluorophore-conjugated secondary antibodies for 1 h at room temperature in the dark. Nuclei were counterstained with DAPI, and coverslips were washed, mounted, and imaged using a Leica SP8 inverted confocal microscope.

### Statistical Analysis

All data are presented as mean ± SD (see figure legends for details of statistical analyses for each dataset) from at least three independent experiments. Statistical analyses were performed using GraphPad Prism software. A p-value ≤ 0.05 was considered statistically significant. *p<0.05, **p<0.01, ***p<0.001, ****p<0.0001.

## Results

### Delayed development and cardiac structural defects in *Prpf8^N1531S^* mutant embryos

We previously identified laterality establishment defects in *Prpf8^N1531S^* homozygous (mutant) embryos. Nearly half of the *Prpf8^N1531S^* mutant embryos showed reversed heart looping, arising from altered node cilia motility (Jiang et al., 2025). Detailed analysis of *Prpf8^N1531S^* mutant embryos at post-cardiac looping stages revealed that these embryos were notably smaller compared to WT or *Prpf8^N1531S^* heterozygous (Het) littermates. Similarly, the hearts of *Prpf8^N1531S^* mutant embryos were also smaller compared with WT hearts (Fig. 1A–D). *Prpf8^N1531S^* mutants exhibit mid-gestation embryonic lethality (Jiang et al., 2025). At E10.5, somite-matched (S31) *Prpf8^N1531S^* mutant hearts, whether displaying correct (CL) or reversed (RL) looping, showed pronounced abnormalities in ventricular trabeculation and compaction zone development (Fig. 1E–H). Both CL and RL *Prpf8^N1531S^* mutant groups exhibited a substantial reduction in ventricular compact zone thickness compared with WT or Het embryos (Fig. 1E). Furthermore, *Prpf8^N1531S^*mutant hearts showed significantly sparser and shorter ventricular trabeculae, compared with the thicker and more extensive trabecular network observed in controls (Fig. 1A–D). Although the total number of trabeculae originating from the compact zone in both ventricles in mutants did not differ significantly from controls (Fig. 1F), the trabecular length was notably reduced in *Prpf8^N1531S^* mutants at E10.5 (Fig. 1G). Quantification of trabecular thickness revealed a significant decrease in both CL and RL *Prpf8^N1531S^* mutants (Fig. 1H). No significant differences were observed between CL and RL *Prpf8^N1531S^* mutant groups for trabecular or compact zone measurements. In comparison with WT controls, *Prpf8^N1531S^* Het embryonic hearts also displayed a significant decrease in compact zone thickness (Fig. 1E), trabecular length (Fig. 1G), and trabecular thickness (Fig. 1H).

**Figure 1.**
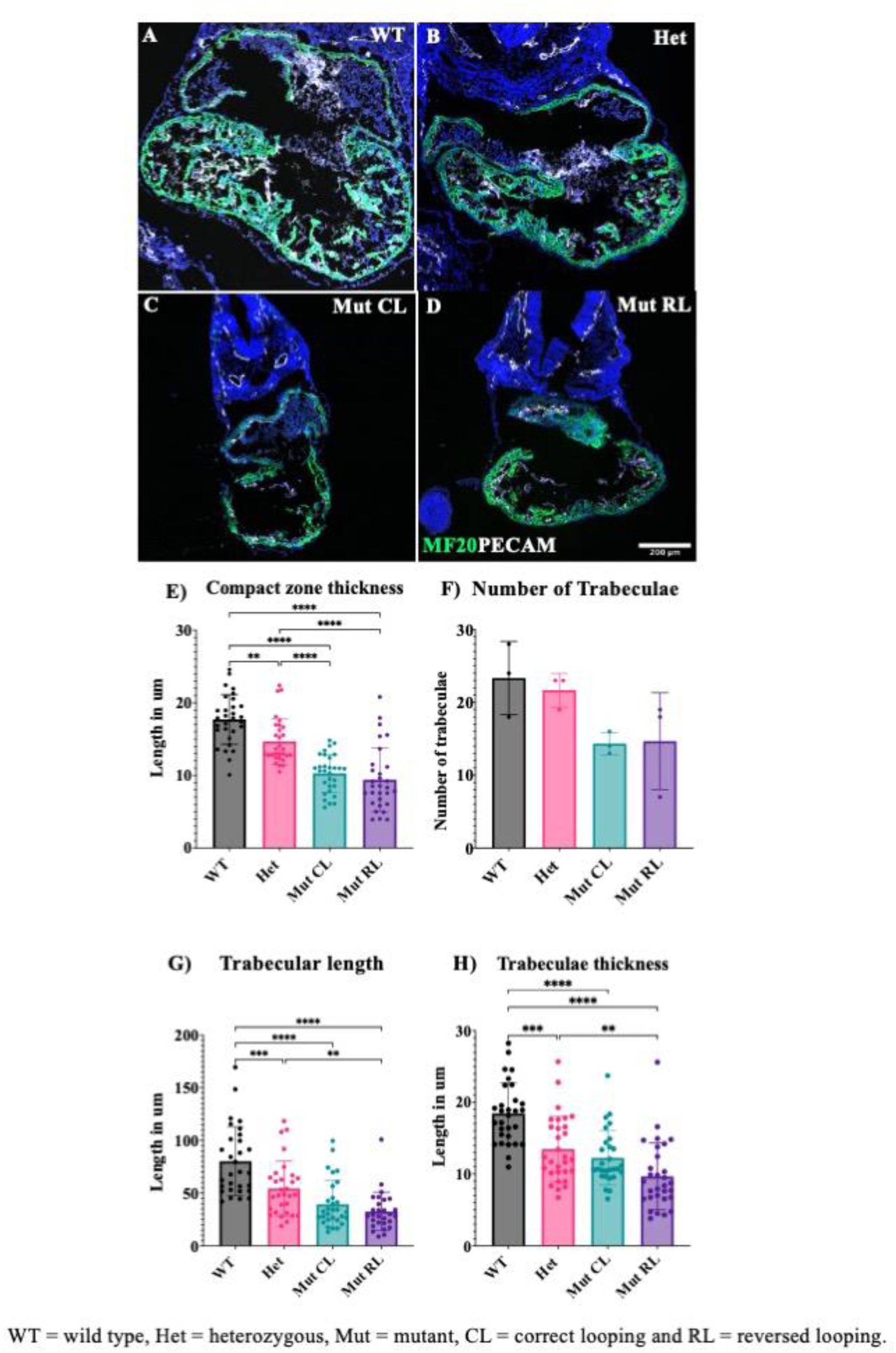
Cardiac morphology analysis in somite-matched (S31) E10.5 wild-type, *Prpf8^N1531S^* heterozygous, and *Prpf8^N1531S^* homozygous (mutant) hearts. CL and RL *Prpf8^N1531S^* mutant hearts were notably smaller compared with WT and Het controls, exhibiting alterations in cardiac trabeculation and compact zone development. A–D) Representative images stained with MF20 (green), PECAM (white), and DAPI (blue) illustrate the cardiac structures of somite-matched (S31) WT, Het, CL, and RL *Prpf8^N1531S^* mutant hearts, respectively. Images were captured at the same magnification. E-H) Quantitative analysis of compact zone thickness, trabecular number, trabecular length, and trabecular thickness, respectively. Data are presented as mean ± SD. Statistical significance was determined by one-way ANOVA with Tukey’s post-hoc test. Scale bar 200 µm. WT = wild type, Het = heterozygous, CL = correct looping and RL = reversed looping.

### Analysis of cell proliferation in stage matched (S31) *Prpf8^N1531S^* mutant hearts

Given the altered ventricular trabeculation and compact zone thickness observed in *Prpf8^N1531S^* mutant hearts, and the established role of cardiomyocyte proliferation in regulating these developmental processes (Chiang et al., 2023, Sandireddy et al., 2019), we assessed cell proliferation in stage-matched E10.5 embryonic hearts from WT, Het, and CL and RL *Prpf8^N1531S^* mutants. The analysis quantified proliferation in endocardial cells, cardiomyocytes, and epicardial cells, as well as overall proliferation present in the left and right ventricles. Immunofluorescence staining of proliferation marker phospho-histone H3 (PH3) alongside cardiac cell markers was used to visualise proliferating cells (Fig. 2A–D). Cardiomyocytes, endocardial cells and epicardial cells showed no significant difference in cell proliferation (Fig. 2E–H). We further analysed the proliferation of cardiomyocytes in the trabeculae and compact zone (CZ) separately (Fig. 2I,J). No significant changes were observed in the percentage of proliferating cells within the trabeculae. However, compact zone cardiomyocytes showed a significant increase in proliferation in RL *Prpf8^N1531S^* mutant hearts compared with WT control. These findings indicate that cardiomyocytes in the compact zone undergo increased proliferation; however, this increase does not contribute to expansion of the trabecular myocardium, suggesting a failure to couple proliferation with trabecular growth.

**Figure 2:**
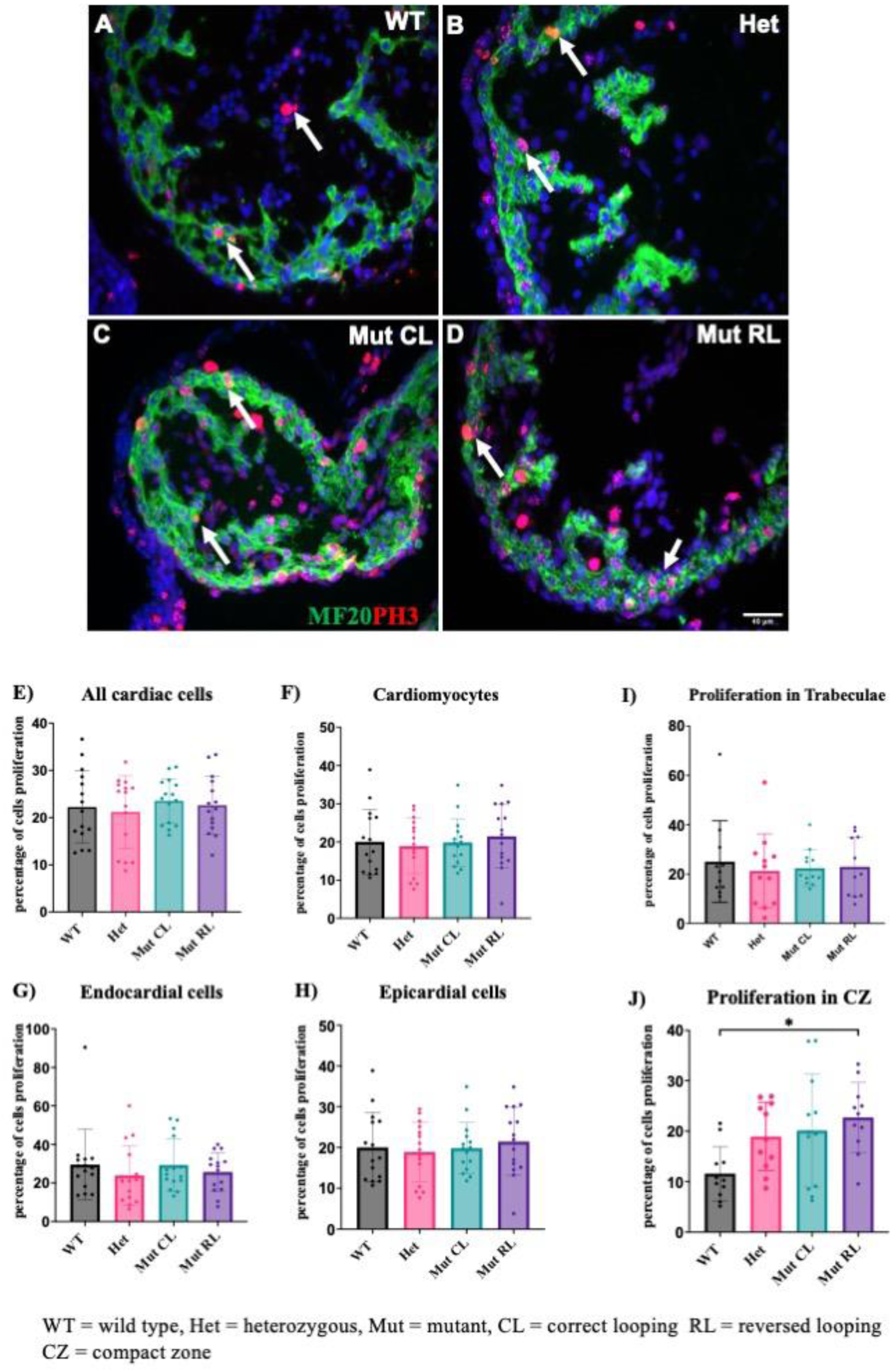
Cell proliferation analysis in somite-matched (S31) cardiac ventricles of E10.5 wild-type, and *Prpf8^N1531S^* homozygous (mutant) hearts. A–D) Representative images of heart sections stained with MF20 (cardiomyocyte marker), PH3 (proliferation marker), and DAPI (nuclear stain). E) Quantification of the percentage of all proliferating cardiac cells in the embryonic heart ventricles, F) percentage of proliferating cardiomyocytes in the ventricles, G) percentage of proliferating endocardial cells in the ventricles, H) percentage of proliferating epicardial cells in the ventricles, I) percentage of proliferating cardiomyocytes in the trabeculae and J) percentage of proliferating cardiomyocytes in the compact zone (CZ). No significant differences in overall cell proliferation were observed. However, a significant increase in proliferating cardiomyocytes (*p* = 0.0108) was detected in the compact zone (CZ) of RL Prpf8^N1531S^ mutant hearts. Data are presented as mean ± SD. Statistical significance was assessed using one-way ANOVA followed by Tukey’s post-hoc test. White arrows indicate proliferating cells. Scale bar, 40 µm. WT = wild type, CL = correct looping, RL = reversed looping.

### Disruption of cellular and cytoskeletal organisation in *Prpf8^N1531S^* mutant hearts

The reduced ventricular trabeculation and compact zone thickness observed in *Prpf8^N1531S^* mutant hearts were not fully accounted for by changes in cardiomyocyte proliferation. Therefore, we next examined whether defects in cardiac marker expression and/or cellular organisation contribute to these morphological differences. Immunofluorescence staining for the sarcomeric myosin protein MF20 in cardiomyocytes and endomucin labelling of endocardial cells revealed notable abnormalities in *Prpf8^N1531S^* mutant hearts (Fig. 3A–C). The cellular arrangement in the compact zones of both CL and RL *Prpf8^N1531S^* mutant hearts appeared random and disorganised, contrasting sharply with the orderly layers observed in WT hearts (Fig. 3A–C). Despite the abnormal cellular organisation observed in *Prpf8^N1531S^* mutant hearts, the cardiomyocyte marker MF20 remained uniformly expressed across all myocardial cells (Fig. 3A–C). Expression analysis of the cardiac transcription factor *Nkx2.5* by *in situ* hybridisation also revealed no alteration in expression pattern (Fig. D–F). However, there was a noticeable disruption in the expression of the endocardial cell marker Endomucin (Fig. 3A–C) and vascular endothelial marker PECAM (Fig. 3G–I) in both CL and RL *Prpf8^N1531S^* mutant hearts, highlighting defects in the endocardial and endothelial cell organisation. The overall morphology of epicardial cells attached to the surface of the heart in *Prpf8^N1531S^*mutants remained unaffected. Immunofluorescence staining for ACTC1 (α-Actinin) showed no evident alteration in sarcomere assembly in CL *Prpf8^N1531S^* mutant hearts. Notably, signs of premature cardiomyocyte maturation, characterised by thick, well-organised sarcomeric bands within embryonic cardiomyocytes (Ahmed et al., 2022), were apparent in RL *Prpf8^N1531S^* mutants (Fig. 3J–L).

**Figure 3:**
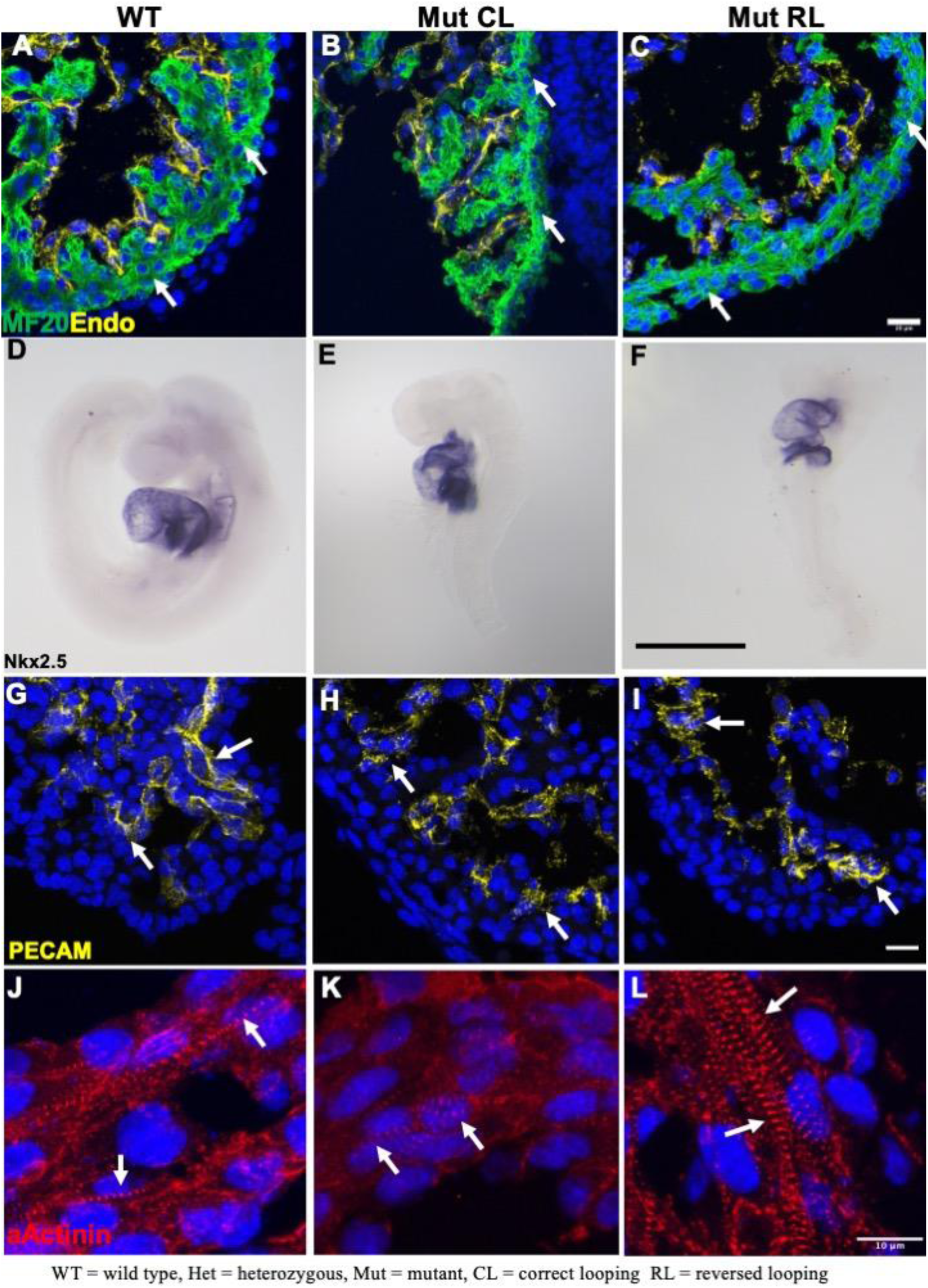
Cardiac and cellular markers expression analysis in somite-matched (S31) cardiac ventricles of E10.5 wild-type, and *Prpf8^N1531S^* homozygous (mutant) hearts. A–C) Cardiomyocyte marker MF20 staining in the mouse embryonic heart. No changes in MF20 staining pattern were observed in CL and RL *Prpf8^N1531S^* mutants. The CL and RL *Prpf8^N1531S^* mutant compact zone cells showed disorganised cellular arrangement compared with WT. D–F) Whole-mount *in situ* hybridisation for *Nkx2.5* in E9.5 embryos. *Nkx2.5* expression in the developing heart is comparable between WT, CL and RL *Prpf8^N1531S^* mutant embryos, indicating preserved early cardiac progenitor specification. G–I) Endocardial cell marker Endomucin staining in the embryonic heart. The CL and RL *Prpf8^N1531S^* mutants showed altered Endomucin localisation compared with control. White arrows indicate the staining pattern. J–L) α-Actinin (ACTC1) staining showing sarcomere organisation in the mouse embryonic heart. The *Prpf8^N1531S^* RL mutant showed thicker sarcomeric striations compared with WT and CL *Prpf8^N1531S^*mutant. Scale bar 20 µm (A–C, G–I), Scale bar 10 µm (J–L), Scale bar 1mm (D–F). WT = wild type, CL = correct looping, RL = reversed looping.

Due to the finding that trabeculae formation in the mouse relies on polarised cell division mediated by cytoskeletal organisation (Grego-Bessa et al., 2007), we sought to investigate whether alterations in the cytoskeletal and membrane-associated structures underlie the trabeculation defects in *Prpf8^N1531S^* mutant hearts. Both CL and RL *Prpf8^N1531S^* mutants displayed disrupted cytoskeletal organisation, suggesting that the *Prpf8^N1531S^* mutation adversely affects the structural integrity of the cytoskeleton. The expression of non-muscle myosin IIB (NM IIB), which is essential for cytoskeletal organisation (Ridge et al., 2017), was altered in *Prpf8^N1531S^* mutant hearts (Fig. 4A–C), especially in RL mutant hearts (Fig. 4C). Phalloidin staining, which labels F-actin (Verderame et al., 1980), also revealed disorganised actin filament arrangement in the *Prpf8^N1531S^* mutant hearts (Fig. 4D–F). β-catenin, crucial for cell-cell adhesion (Gottardi and Gumbiner, 2001), displayed aberrant expression patterns at the cell membrane in *Prpf8^N1531S^* mutant hearts (Fig. 4G–I). WGA staining in *Prpf8^N1531S^* mutant hearts further revealed irregular cell membrane organisation (Fig. 4J–L), with RL mutants exhibiting severe disarray and poorly defined cell boundaries (Fig. 4L). Collectively, these findings illustrate that the *Prpf8^N1531S^* mutation leads to profound disruptions in cell arrangement, cytoskeletal organisation and cell membrane integrity in embryonic hearts.

**Figure 4:**
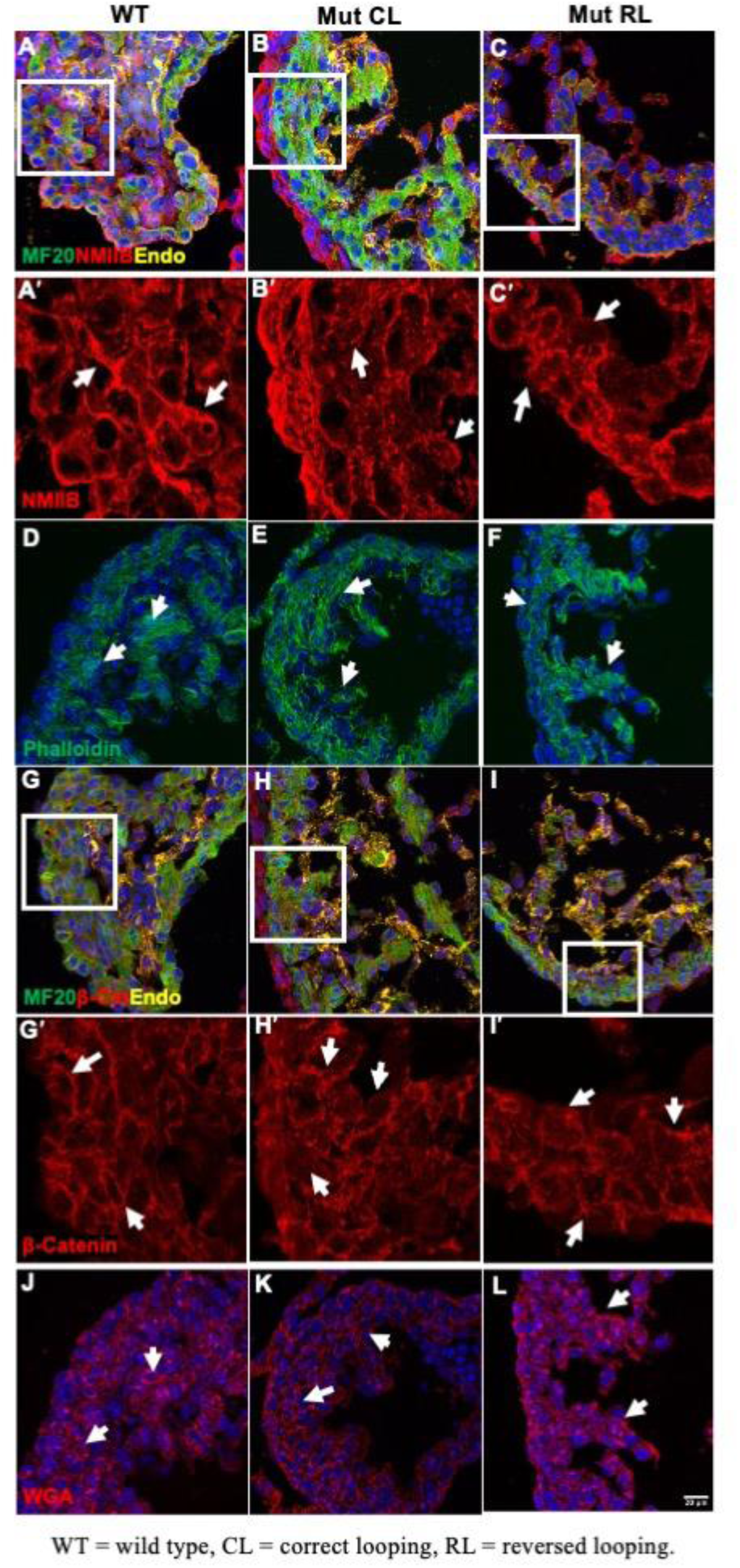
Cytoskeleton markers expression analysis in somite-matched (S31) cardiac ventricles of E10.5 wild-type, and *Prpf8^N1531S^* homozygous (mutant) hearts. A–C, A′–C′) NM II-B staining showing cytoskeletal organisation in the mouse embryonic heart. Images A′–C′ show zoomed NM II-B staining only corresponding to panels A–C. The *Prpf8^N1531S^* CL and RL mutants showed a disturbed expression pattern compared to control. D–F) Phalloidin staining showing the F-actin organisation in the mouse embryonic heart. The *Prpf8^N1531S^* CL and RL mutants showed altered phalloidin staining pattern compared with control. G–I, G′–I′) β-catenin staining showing cell membrane organisation in the mouse embryonic heart. Images G′–I′ show zoomed β-catenin staining corresponding to panels G–I. The *Prpf8^N1531S^* CL and RL mutants showed a disturbed and discontinued β-Catenin expression at the cell membrane. J–L) WGA staining showing cell membrane organisation in the mouse embryonic heart. The *Prpf8^N1531S^* CL and RL mutants showed a disturbed and discontinued WGA expression at the cell membrane. White arrows indicate the staining pattern. Scale bar 20µm. WT = wild type, CL = correct looping, RL = reversed looping.

### Disturbed extracellular matrix protein expression in *Prpf8^N1531S^* mutant hearts

Mouse mutants with trabeculation defects display a reduction in extracellular matrix (ECM) composition during cardiac development (Grego-Bessa et al., 2023), as cytoskeletal integrity directly influences cell–ECM interactions during heart development (Israeli-Rosenberg et al., 2014, Silva et al., 2021). To evaluate ECM production in *Prpf8^N1531S^*mutants, we performed immunofluorescence staining using antibodies against key ECM components. Fibronectin 1 (FN1), which is secreted into the extracellular space, showed altered protein distribution between WT and *Prpf8^N1531S^*mutant hearts (Fig. 5A–C, A’–C’). In WT hearts, FN1 displayed a consistent and organised distribution throughout the heart section (Fig. 5A, A’). However, in CL and RL *Prpf8^N1531S^* mutant hearts, FN1 expression was significantly disrupted, especially in the compact zone of RL mutant hearts (Fig. 5B, B’, C, C’), indicating altered ECM organisation. The basement membrane ECM protein Collagen IV (Col IV) staining demonstrated a marked reduction in the mutant hearts compared with WT (Fig. 5D–F, D’–F’). The WT samples exhibited a robust network of Col IV, whereas CL *Prpf8^N1531S^*mutant hearts showed irregular basement lining, along with altered deposition in extracellular spaces. This pattern further intensified in RL *Prpf8^N1531S^*mutant hearts with no Col IV basement marking of cellular structures along with altered extracellular spaces deposition compared to WT (Fig. 5D’–F’). Another basement membrane ECM protein, Laminin (LAM), also exhibited detectable differences between genotypes (Fig. 5G–I, G’–I’). WT hearts displayed a well-defined Laminin matrix surrounding trabeculae and compact zone structures. In contrast, both CL and RL *Prpf8^N1531S^*mutant hearts showed a fragmented and patchy Laminin distribution, especially in RL *Prpf8^N1531S^* mutant hearts (Fig. 5G’–I’). This suggests potential basement membrane defects in the *Prpf8^N1531S^* mutant hearts. Another ECM marker, Hyaluronan Synthase 2 (HAS2), showed notably reduced expression in the mutant hearts compared to WT (Fig. 5J–L, J’–L’). WT samples showed a uniform HAS2 distribution around the cell nucleus. Mutant hearts, however, exhibited a sparse and uneven HAS2 pattern (Fig. 5J–L), suggesting impaired synthesis and maintenance of hyaluronan. HAS2 is an enzyme responsible for the synthesis of hyaluronan, a major component of the ECM, and has a reported role in cardiac trabeculation and compaction (Camenisch et al., 2000).

**Figure 5:**
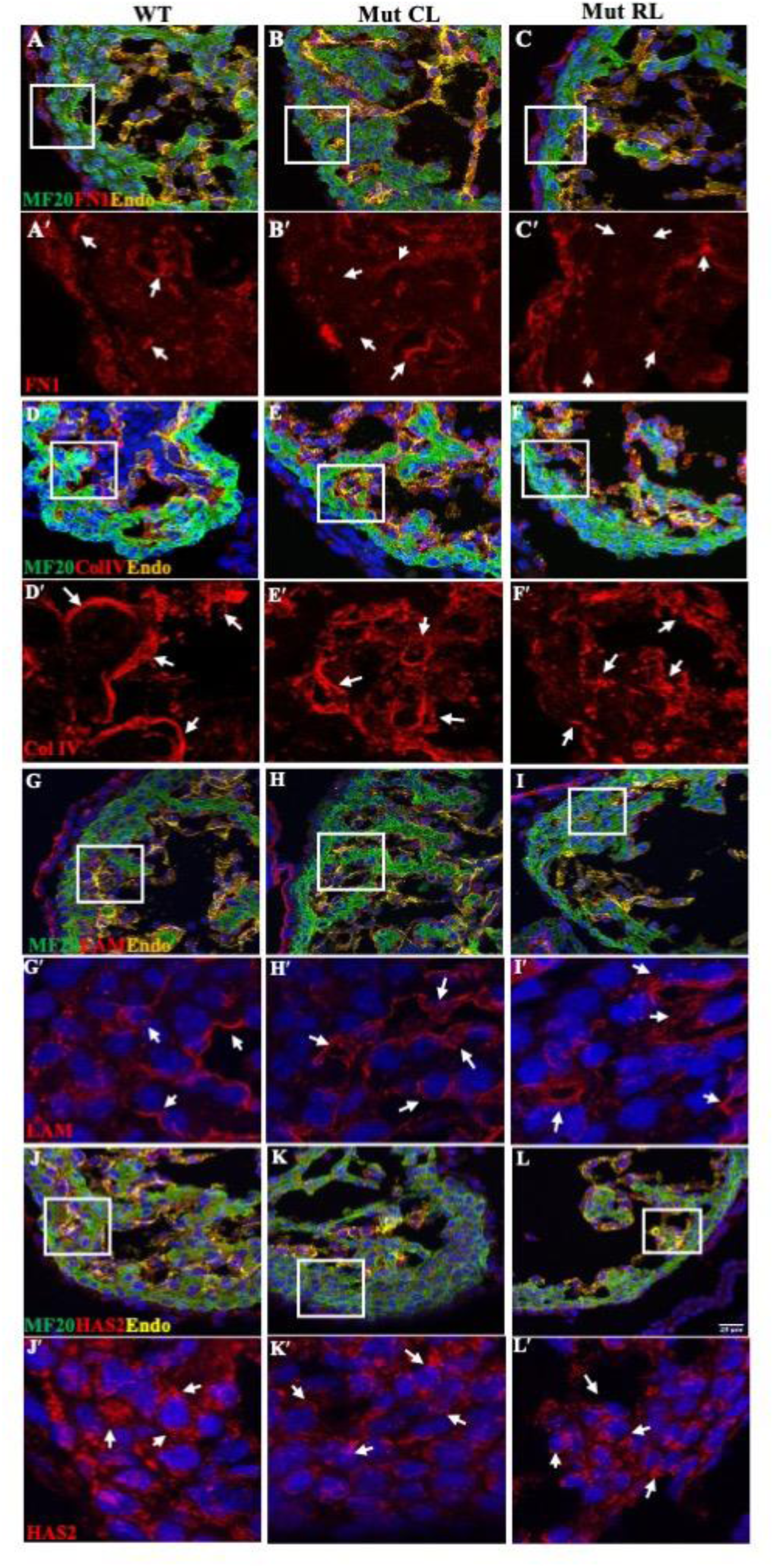
Extracellular matrix protein expression analysis in somite-matched (S31) cardiac ventricles of E10.5 wild-type, and *Prpf8^N1531S^*homozygous (mutant) hearts. A–C, A′–C′) Fibronectin 1 (FN1) staining in the mouse embryonic heart. Images A′–C′ show zoomed FN1 staining corresponding to panels A–C. The ECM protein FN1 is secreted into the extracellular space, and its expression is altered and reduced in *Prpf8^N1531S^* mutant hearts. D–F, D′–F′) Collagen IV (Col IV) staining in mouse embryonic hearts. Images D′–F′ show zoomed Col IV staining corresponding to panels D–F. Col IV, a basement membrane protein, shows altered deposition, particularly in RL *Prpf8^N1531S^*mutant hearts. G–I, G′–I′) Laminin (LAM) staining in the mouse embryonic heart. Images G′–I′ show zoomed LAM staining corresponding to panels G–I. LAM, a basement membrane protein, shows altered expression, particularly in RL *Prpf8^N1531S^*mutant hearts. J–L, J′–L′) Hyaluronan Synthase 2 (HAS2) staining, an enzyme responsible for the synthesis of hyaluronan, a major component of the ECM. Images J′–L′ show zoomed HAS2 staining corresponding to panels J–L. *Prpf8^N1531S^*mutant hearts showed reduced expression in the extracellular space surrounding cells. White arrows indicate ECM protein expression patterns. Scale bar, 20 µm. WT = wild type, CL = correct looping, RL = reversed looping.

### Identification of *Tead1* mis-splicing in *Prpf8^N1531S^* mutant embryos

To determine if splicing abnormalities caused by the *Prpf8^N1531S^* mutation could directly result in the cardiac abnormalities present in mutant embryos, we assessed PRPF8 expression within the developing heart. PRPF8 protein was predominantly confined to the cell nucleus and remained unchanged in both CL and RL *Prpf8^N1531S^* mutant hearts compared with WT controls (Fig. 6A–C). Moreover, cardiomyocytes within both the compact zone and trabecular myocardium of E10.5 *Prpf8^N1531S^* mutant hearts exhibited PRPF8 staining patterns comparable to those observed in WT hearts. These findings indicate that the PRPF8 N1531S mutation does not alter PRPF8 abundance, stability or localisation in the developing heart, suggesting that the cardiac phenotype may instead arise from defects in RNA splicing of key cardiac regulators. Bulk RNA sequencing (RNA-seq) analysis of E10.5 WT, CL and RL Prpf8N1531S mutant embryos (Jiang et al., 2025) revealed aberrant splicing of the cardiac transcription factor Tead1. Given the associated trabeculation defects and the established role of TEAD1 in myocardial wall development (Wen et al., 2019, Liu et al., 2017),. we further investigated its splicing using rMATS, which identified significant exon-skipping events. Specifically, exon 8 skipping was detected in CL mutants (FDR = 6.72 × 10⁻⁴), while exon 5 skipping was preferentially observed in RL mutants (FDR = 1.30 × 10⁻⁵) compared to WT controls. Although not the most statistically significant event genome-wide, Tead1 was prioritised based on its functional relevance to cardiac development.

**Figure 6:**
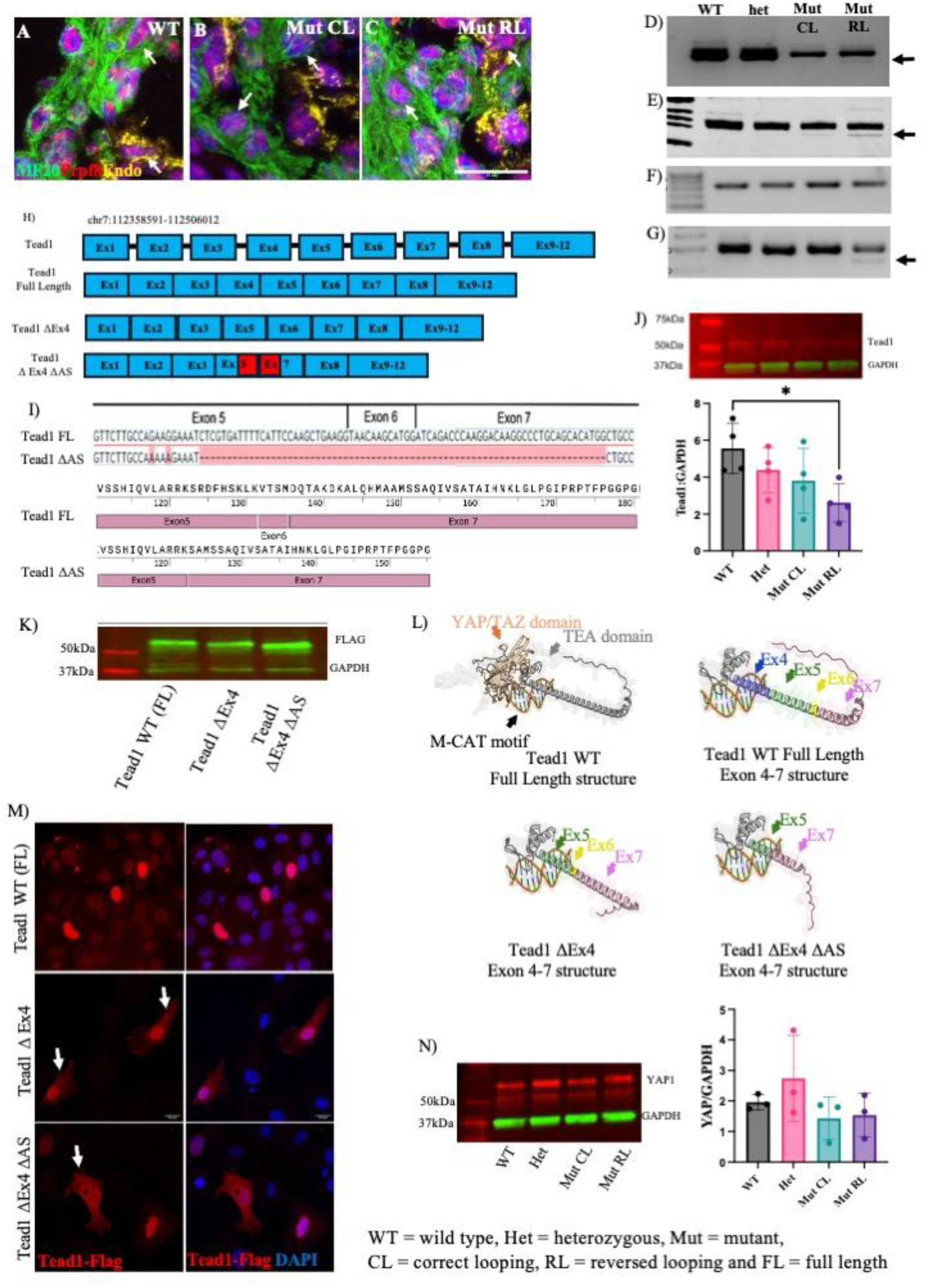
Analysis of *Tead1* mis-splicing in *Prpf8^N1531S^* mutant embryos. A–C) PRPF8 staining in mouse embryonic heart. The CL and RL *Prpf8^N1531S^*mutant hearts showed no change in PRPF8 nuclear localisation pattern compared with WT. D) Agarose gel image of RT-PCR (exon 2-10) of *Tead1* isoforms in cardiac tissues in E10.5 embryos. Arrow indicating the fainter smaller size band in RL *Prpf8^N1531S^* mutant. E) 8% Polyacrylamide gel image of RT-PCR (using primers in exon 2 and 10) of *Tead1* isoforms in E10.5 cardiac tissues, F) Agarose gel image of RT-PCR (using primers in exons 2 and 3) of *Tead1* isoforms in cardiac tissues, G) Agarose gel image of RT-PCR (exon 3-7) of *Tead1* isoforms in cardiac tissues, H) Schematic representation of the mouse *Tead1* exons, showing the full-length transcript (*Tead1*-FL), the exon 4-deleted isoform (*Tead1*-ΔEx4), and the isoform lacking both exon 4 and alternative splicing region which creates a fusion between exon 5 and exon 7 (*Tead1*-ΔEx4 ΔAS). The red box indicates the deleted regions within exons 5 and 7 that generate this fusion. I) *Tead1* full length (FL) and alternate spliced ΔAS product sequence alignment to mouse transcriptome (GRCm39/mm39), J) Western blot image of TEAD1 protein in E10.5 whole embryos lysate (TEAD1 staining shown in red; GAPDH shown in green). Western blot image quantification of TEAD1 protein expression relative to GAPDH showed significant downregulation in RL *Prpf8^N1531S^* mutant, K) Western blot image of TEAD1 epitope-tagged isoforms expressed in HEK cells. The isoforms with exon deletions produce proteins of smaller molecular weight. L) The structures of full-length WT and alternatively spliced TEAD1 isoforms were predicted using AlphaFold3 and subsequently visualised in PyMOL, with DNA modelled based on the M-CAT sequence (Holden and Cunningham, 2018a). TEAD1 WT full length structure of region Exon 4-7; Tead1 Δ Ex4 structure of exon 4-7, showing missing exon 4; and Tead1 Δ Ex4 ΔAS structure of Exon 4-7. Showing missing Δ Ex4 and AS region. M) Immunofluorescence image of H9c2 cell transfected with flagged tagged TEAD1 WT- full length (FL), TEAD1 Δ Ex4 and TEAD1 ΔEx4 Δ AS isoforms. The WT showed showed nuclear localisation, while some cells in the mis-spliced isoforms showed localisation in cytoplasm and nucleus. Arrows indicating cytoplasmic localisation of flagged tagged TEAD1. N) Western blot image of YAP1 protein in E10.5 whole embryos lysate. The bands quantification of YAP1 protein expression relative to GAPDH shows no significant change in *Prpf8^N1531S^* mutant. Data are presented as mean ± SD. Statistical significance was assessed using one-way ANOVA with Tukey’s post-hoc test. Scale bar 20 µm. WT = wild type, Het = heterozygous, Mut = mutant, CL = correct looping, RL = reversed looping and FL = full length

We designed primers to amplify the *Tead1* sequence between exons 2–10 to confirm exon 5 and 8 skipping in the *Prpf8^N1531S^* mutant embryonic hearts (Fig. 6D). Both WT and *Prpf8^N1531S^*het E10.5 embryonic heart cDNA produced multiple PCR products on an agarose gel, confirming alternative splicing of the *Tead1* transcript consistent with annotations of the reference mouse transcriptome (GRCm39/mm39) (Fig 6D). However, in CL and RL *Prpf8^N1531S^* mutants, several bands were either faint or absent on agarose gel electrophoresis (Fig. 6D). A very faint band of lower size was present in cDNA isolated from RL *Prpf8^N1531S^* mutant embryonic hearts (Fig. 6D indicated with arrow). We repeated the same experiment using a polyacrylamide gel to separate putative splice variants at higher resolution, confirming that this smaller, fainter band is present in both CL and RL *Prpf8^N1531S^*mutants (Fig. 6E). RT-PCR using primer pairs to amplify exon 2-3 (Fig. 6F) and exon 3-7 (Fig. 6G) of *Tead1* further verified the location of this mis-spliced product between exon 3 to 7. This shorter mis-spliced RT-PCR product in RL *Prpf8^N1531S^* mutant was sequenced, revealing a novel alternatively spliced *Tead1* transcript. This isoform lacks exon 4 (ENSMUST00000059768.18 Chr7: 112,441,095-112,441,157;), part of exon 5, the complete exon 6, and part of exon 7 were missing (ENSMUST00000059768.18 Chr7: 112,441,284-112,456,041; Fig. 6H and I). This mis-spliced transcript was not detected in our RNA-seq data and is not annotated as a known *Tead1* isoform. We therefore refer to this variant as *Tead1*ΔEx4 ΔAS.

We further verified the sequence of the *Tead1* PCR product using TA cloning of individual products from the RT-PCR reaction. The RT-PCR produced multiple *Tead1* products with a combination of skipped annotated exons; the exon 4 and 6 skipping events were present in multiple known transcripts. Along with these known *Tead1* alternate transcripts, we identified the presence of alternatively spliced *Tead1* transcript *Tead1* ΔEx4 ΔAS as identified earlier. This *Tead1* ΔEx4 ΔAS product results in the deletion of exon4 and another 25 amino acid sequence from TEAD1 protein (Serine 123 to Methionine 147), with Alanine 148 being substituted by Serine 123, thereby maintaining the reading frame (Fig 6I).

The *Prpf8^N1531S^* mutant RNA-seq data (Jiang et al., 2025) also showed significantly altered differential gene expression (DE fold change = -1.1154332, DE Padj 1.28 x 10^-5^) of *Tead1* only in RL *Prpf8^N1531S^* mutant embryos. Western blot analysis of Tead1 protein using E10.5 whole embryo lysate showed a trend towards downregulation of protein levels in Het, CL and RL *Prpf8^N1531S^*mutant embryo compared to WT control, with the RL *Prpf8^N1531S^* mutant showing a significant difference (Fig. 6J).

We further analysed the effect of the *Tead1* ΔAS transcript alterations on protein expression. We generated N-terminal FLAG tagged mammalian cell expression plasmids containing mouse Tead1 full length (Tead1-FL), Tead1 without exon 4 (Tead1 ΔEx4), and Tead1 without both exon 4 and the alternatively spliced exons 5-7 (Tead1 ΔEx4 ΔAS) (the alternatively spliced transcript lacks exon 4 as well). These constructs were transfected into human HEK293 cells. As expected, an anti-FLAG western blot performed on extracts from the transfected cells showed lower molecular weight proteins generated from transfection of Tead1 ΔEx4 and Tead1 ΔEx4 ΔAS plasmids compared to Tead1-FL, confirming that a stable protein of reduced molecular weight is generated from the TeadΔEx4 and Tead1 ΔEx4 ΔAS transcript (Fig. 6K). In silico structural modelling of mutant TEAD1 isoforms using AlphaFold3 suggests that deletion of Ex4 and the ΔAS region may perturb the TEA DNA-binding domain and adjacent linker region, which is important for nuclear localisation (Magico and Bell, 2011, Calses et al., 2023). The model also predicts that the TEAD1 YAP/TAZ domain structure remains intact (Fig. 6L). To investigate the role of the mis-spliced region of TEAD1 in nuclear localisation, we transfected WT (full length) and mis-spliced *Tead1* variants into the rat cardiomyocyte H9c2 cell line. Cells transfected with WT *Tead1* showed predominantly nuclear localisation. In contrast, *Tead1* mis-spliced variants exhibited mixed nuclear and cytoplasmic expression in a subset of cells (Fig. 6M), with a pronounced accumulation around the nuclear lamina, forming a ring-like structure. These aberrant localisation patterns were further enhanced in cells transfected with the Tead1 ΔEx4ΔAS variant, confirming *Tead1* mis-splicing affects TEAD1 nuclear translocation or localisation. Which may result in altered Tead1 transcription factor down stream targets activation.

To investigate the impact of *Tead1* mis-splicing on the Hippo pathway, which regulates organ size, we measured the expression levels of Yes-associated protein 1 (YAP1) in both control and *Prpf8^N1531S^* mutant embryos. YAP1 serves as a key transcriptional regulator within the Hippo signalling pathway, interacting with TEA domain (TEAD) transcription factors in the nucleus. This interaction activates downstream signalling pathways associated with TEAD, influencing cell proliferation and growth (Fu et al., 2022, Chen et al., 2010). Our analysis showed no significant difference in YAP1 expression in WT, Het and CL and RL *Prpf8^N1531S^* mutant embryos (Fig. 6N), suggesting that the defects observed in *Prpf8^N1531S^* mutant are due to *Tead1* mis-splicing affecting the structure of the TEA DNA-binding domain and its adjacent linker segment, but not reducing TEAD1 YAP/TAZ domain protein interactions.

### Tead1 target gene expression is altered in *Prpf8^N1531S^* mutant embryos

Given that TEAD1 regulates transcriptional programs essential for myocardial–endocardial signalling during early cardiac morphogenesis, we next examined whether *Tead1* mis-splicing in *Prpf8^N1531S^* mutant embryos affected the Notch signalling pathway, which is a key driver of endocardial activation and trabeculation (Grego-Bessa et al., 2007). The *Prpf8^N1531S^* mutant RNA-seq data identified the downregulation of Notch signalling pathway genes including *Hey2*, *Hey1*, and *Heyl* which are important for cardiac trabeculation (Grego-Bessa et al., 2007). Therefore, stage matched E10.5 control and *Prpf8^N1531S^* mutant hearts were stained for the intracellular domain of the NOTCH1 receptor (NICD1) (Fig. 7A) to identify activated NOTCH signalling. NICD1 expression was downregulated in mutant endocardial cells, with RL *Prpf8^N1531S^* mutants hearts showing a significant difference compared to controls (Fig 7A). We further confirmed the downregulation of NICD1 expression in the E10.5 *Prpf8^N1531S^* mutant whole embryo lysate by Western blot. The RL *Prpf8^N1531S^* mutants showed significant NICD1 downregulation compared to WT embryos (Fig. 7B).

**Figure 7:**
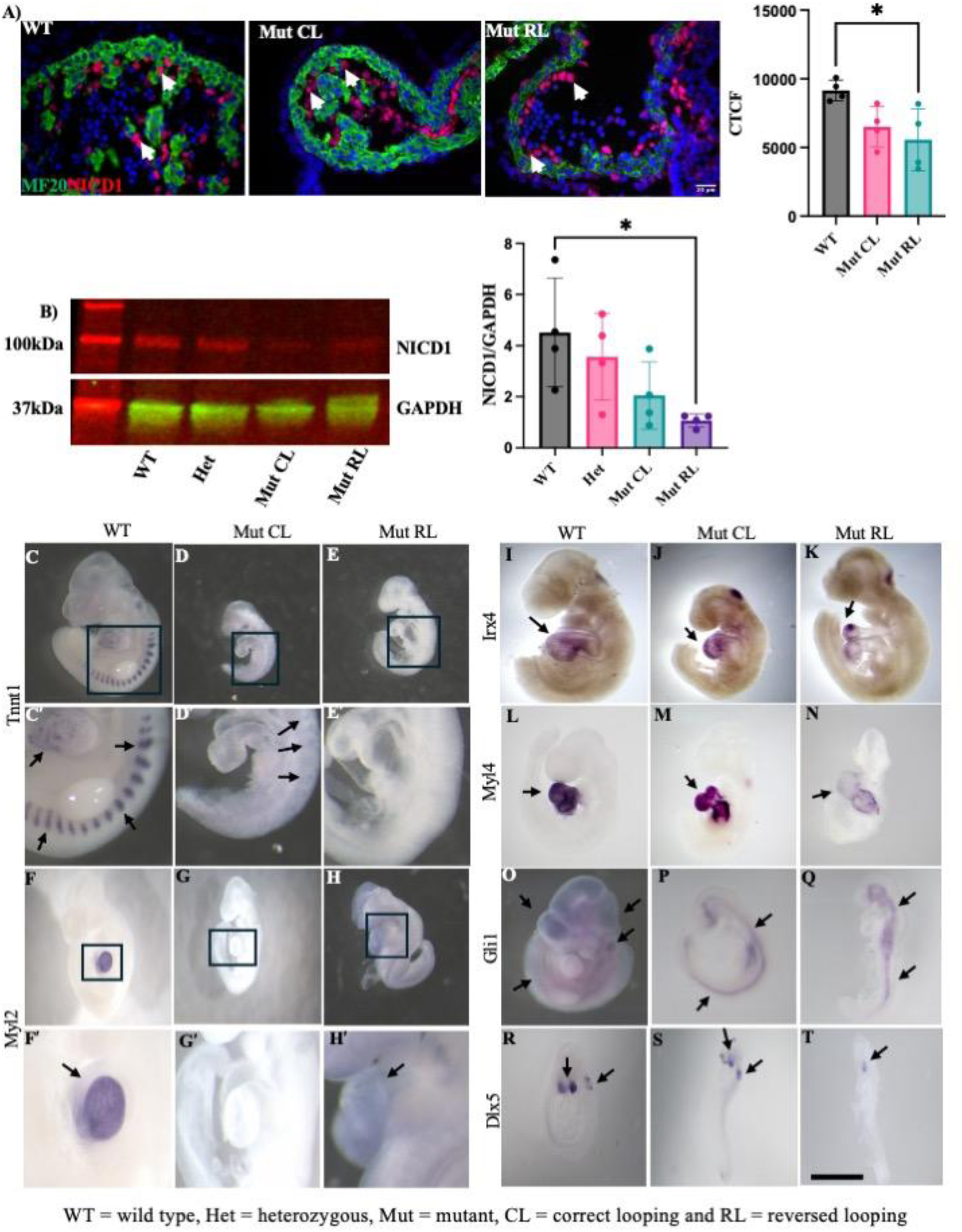
Dysregulation of TEAD1 target genes and reduced Notch signalling in *Prpf8^N1531S^* mutant embryos. A) Representative images of E10.5 heart sections from WT, CL and RL *Prpf8^N1531S^*Mutants stained with NICD1. Quantification of NICD1 expression in endocardial cells within the cardiac ventricles shows significantly (*p*=0.0301) reduced expression in RL *Prpf8^N1531S^* mutant hearts. (CTCF=Corrected Total Cell Fluorescence). B) Representative western blot image showing reduced NICD1 levels (∼100 kDa) in *Prpf8^N1531S^* mutant embryos. GAPDH (∼37 kDa) serves as a loading control. Quantification of NICD1 protein levels normalised to GAPDH shows significant downregulation in RL *Prpf8^N1531S^* mutant embryos (*p*=0.0323) compared with WT control. Data are presented as mean ± SD. Statistical significance was assessed using one-way ANOVA with Tukey’s post-hoc test. Scale bar 20 µm. C-E, C′-E′) Whole-mount *in situ* hybridisation showing expression of the sarcomeric gene *Tnnt1* in E10.5 embryos. Compared with robust expression in WT embryos, *Tnnt1* expression is reduced in CL *Prpf8^N1531S^* mutants and nearly absent in RL *Prpf8^N1531S^*mutants. Images C′–E′ show zoomed *Tnnt1* staining corresponding to panels C-E. F-H) *In situ* hybridisation for *Myl2*, a ventricular cardiomyocyte marker, reveals strong ventricular-specific expression in WT embryos, with markedly reduced or absent expression in CL and RL *Prpf8^N1531S^*mutant embryos. Images F′–H′ show zoomed *Myl2* staining corresponding to panels F-H. I-K) Expression of the ventricular identity gene *Irx4* is reduced in CL *Prpf8^N1531S^* mutant embryos and further diminished in RL mutant embryos compared with WT controls. L-N) *In situ* hybridisation for *Myl4* shows reduced myocardial expression in both Mut CL and Mut RL embryos relative to WT. O-Q) Expression of the Hedgehog pathway target *Gli1*, which is decreased in CL *Prpf8^N1531S^* embryos and further reduced in RL *Prpf8^N1531S^* mutant embryos, with residual expression confined to midline regions. R-T) *In situ* hybridisation for the neural crest associated gene *Dlx5* at E9.5 shows reduced expression in CL *Prpf8^N1531S^* mutant embryos and near-complete loss in RL *Prpf8^N1531S^* mutant embryos compared with WT. Scale bar 1mm (C-T). WT = wild type, CL = correct looping and RL = reversed looping.

A previous study identified cardiac sarcomeric genes regulated by TEAD-dependent transcriptional programmes (Akerberg et al., 2019). We, therefore, examined the expression of several of these genes in the CL and RL *Prpf8^N1531S^*mutant embryos to check for expression abnormalities that may arise from the altered TEAD1 protein isoform predicted in CL and RL *Prpf8^N1531S^* mutant embryos. Consistent with the variation in developmental defects observed in the *Prpf8^N1531S^* mutant embryos, the expression pattern of TEAD1 downstream targets also varies between *Prpf8^N1531S^* mutant samples. In E10.5 WT embryos, *Tnnt1* (Troponin T1) gene expression was observed in the heart and in the somite region (Fig. 7C–E, C′-E′). Compared to WT, in CL *Prpf8^N1531S^*embryos the *Tnnt1* expression was absent from the heart, but a fainter expression was seen in the somite region (Fig. 7D, D′). However, the RL *Prpf8^N1531S^* mutants showed a complete disappearance of the *Tnnt1* expression in the embryo (Fig. 7E, E′). Another target gene *Myl2* (Myosin Light Chain 2) showed expression within the cardiac ventricle region, which was either fainter or completely missing in the CL and RL *Prpf8^N1531S^* mutants (Fig. 7F–H, F′-H′). Furthermore, the Tead1 target *Irx4* (Iroquois Homeobox 4), which is expressed in the cardiac ventricle region, also showed reduced expression in mutants, especially in the RL *Prpf8^N1531S^* heart (Fig. 7I–K). The sarcomeric structural gene *Myl4* also showed reduced expression in *Prpf8^N1531S^* mutant hearts (Fig. 7L–N).

Akerberg et al, (2019) also identified genes downregulated in TEAD1 knockout mouse embryos. We sought to determine if the expression of some of these genes was altered in *Prpf8^N1531S^* mutant embryos. *Gli1* gene expression was downregulated in CL and RL *Prpf8^N1531S^* E9.5 embryos compare to WT. The expression was mainly confined to the midline region of mutant embryos, which was further decreased in RL *Prpf8^N1531S^* mutant embryos (Fig. 7O–Q). *Dlx5*, which is expressed in neural crest cells, also showed decreased expression in *Prpf8^N1531S^*mutants compared to WT. In E9.5 WT embryos, *Dlx5* expression can be seen in rhombomeres, pharngyeal areches and in the craniofacial region. However, this expression was reduced in CL mutants, with the expression almost missing in the rhombomeres and craniofacial region in RL *Prpf8^N1531S^* mutant (Fig. 7R–T). Together, these results indicate that Tead1 mis-splicing in *Prpf8^N1531^* mutant embryos disrupts TEAD1-dependent gene regulation and Notch signalling, contributing to the cardiac developmental abnormalities.

## Discussion

This study identifies *Prpf8* as a previously unrecognised regulator of ventricular wall morphogenesis and links a hypomorphic spliceosomal defect to selective disruption of developmentally critical cardiac transcriptional and signalling programmes rather than global splicing failure. As a core catalytic component of the spliceosome (Malinová et al., 2017), partial loss of PRPF8 function is likely to preferentially affect splicing of a subset of transcripts. In *Prpf8^N1531S^* mutants, we propose that the affected transcripts include those required for cardiac trabeculation and compaction (Sedmera et al., 2000, Grego-Bessa et al., 2007, Zhang et al., 2013, Meyer et al., 2020).

The *Prpf8^N1531S^* mutant embryos exhibit mis-splicing of multiple developmentally important transcripts, including the ciliary small GTPase *Arl13b* (Jiang et al., 2025). However, mis-splicing in *Prpf8^N1531S^* mutants also affects multiple non-ciliary genes, suggesting that cardiac ventricular wall abnormalities detected in the mutants may arise from combined disruption of ciliary and non-ciliary pathways. The central finding of this study is the aberrant splicing of the transcription factor *Tead1* in *Prpf8^N1531S^*mutant embryos, generating an in-frame isoform with reduced abundance and widespread dysregulation of TEAD1-associated gene expression. TEAD1 is a major downstream effector of the Hippo signalling pathway (Mia and Singh, 2019). Through its integration of mechanical and transcriptional cues via the Hippo–YAP/TAZ–TEAD pathway, TEAD1 regulates cardiomyocyte proliferation, differentiation, and ventricular wall development (Yamada et al., 2023, Liu et al., 2020, Liu et al., 2019, Chen et al., 1994, Liu et al., 2017). TEAD1 contains a conserved TEA DNA-binding domain that recognises M-CAT motifs in promoters and enhancers, and a C-terminal transactivation domain that engages cofactors such as YAP, TAZ, and VGLL proteins to drive transcription. These domains are linked by a region important for protein stability and nuclear localisation (Joshi et al., 2017, Li et al., 2022, Yoshida, 2008b, Holden and Cunningham, 2018b, Yoshida, 2008a, Jiang et al., 2000, Pobbati et al., 2012, Calses et al., 2023).

The alternatively spliced *Tead1* (Tead1 ΔAS) transcript identified in *Prpf8*^N1531S^ mutant embryos removes residues immediately C-terminal to the TEA DNA-binding domain, within the linker region. In *Drosophila*, the TEAD homolog *Scalloped* contains a functional nuclear localisation signal located just downstream of the TEA domain, and disruption of this region results in TEAD cytoplasmic retention and reduced transcriptional activity (Magico and Bell, 2011). Studies in mammalian systems demonstrate that mutation or deletion of conserved basic residues outside the core TEA domain impairs nuclear localisation of TEAD proteins, indicating that efficient nuclear import depends on sequences spanning the TEA-linker interface (Calses et al., 2023). Therefore, deletion of exon 4 and the Ser¹²³-Met¹⁴⁷ region in the *Tead1* ΔAS isoform likely reduces nuclear accumulation of TEAD1 as observed in H9C2 cells (Figure 6M), limiting the availability of transcriptionally active TEAD1 in the nucleus and attenuating TEAD1-dependent gene regulation during cardiac development.

Two independent studies demonstrated that TEAD1 splicing modulates Hippo-YAP signalling through alternative inclusion of the 12-nucleotide exon 6 (Verma et al., 2022, Isaac et al., 2024), generating isoforms with altered transcriptional activity and YAP responsiveness. Although TEAD1 transcriptional output is commonly regulated by Hippo pathway control of YAP1, total YAP1 protein levels were unchanged in *Prpf8^N1531S^*mutant embryos, consistent with the absence of predicted structural alterations in the TEAD1 YAP/TAZ-binding domain. This finding suggests that the observed TEAD-associated transcriptional dysregulation may arise from altered TEAD1 isoform composition and/or DNA-binding competence, not from changes in YAP1 levels. These observations support a model in which isoform dosage and context, rather than exon loss alone, determine TEAD1 activity during cardiac development. An intriguing observation in the phenotype is the increase in cardiomyocyte proliferation within the compact zone despite a reduction in compact zone thickness, indicating that ventricular wall growth is not simply determined by cell-cycle entry. Trabeculation and compaction require coordinated tissue architecture, cytoskeletal tension, cell-cell adhesion, and ECM organisation (Wu, 2018), all of which are disrupted in *Prpf8^N1531S^* mutant hearts. TEAD1 integrates Hippo-mechanical cues into transcriptional programs that regulate extracellular matrix composition, cell-matrix adhesion, and cytoskeletal tension (Dupont et al., 2011, Song et al., 2024, Pagliari et al., 2021). In this context, elevated proliferation may reflect a compensatory response to impaired morphogenesis or differentiation, rather than productive compact zone expansion and directional cardiomyocyte migration into trabeculae (Li et al., 2016).

The finding of reduced NICD1 and attenuated Notch pathway readouts (*Hey1, Hey2, Heyl* (Jiang et al., 2025)) in *Prpf8^N1531S^* mutants is particularly important because endocardial Notch signalling acts as a gatekeeper of ventricular trabeculation (Grego-Bessa et al., 2007). TEAD1 dysfunction caused by mis-splicing and reduced transcriptional activity may have perturbed myocardial-endocardial coupling, thereby directly or indirectly attenuating NOTCH signalling activation in the trabecular endocardium. Supporting this concept, previous work has shown that YAP-TEAD transcriptional activity can up-regulate the Notch ligand *Jagged1*, leading to activation of NOTCH signalling (Tschaharganeh et al., 2013). This suggests that reduced TEAD1 activity may compromise the myocardial competence to sustain endocardial Notch signalling. Such a mechanism would combine several of our observations, including diminished trabecular thickness and length despite preserved trabeculation initiation, altered endocardial integrity, and the downstream loss of trabeculation-supporting cues.

*Tead1* mis-splicing and loss of function in *Prpf8^N1531S^*mutant embryos is likely the cause of *Tead1* target gene mis-expression (Akerberg et al., 2019). Notably, sarcomeric genes such as *Tnnt1*, *Myl2*, and *Myl4* are known to be regulated by TEAD-dependent transcriptional programs involved in cardiac muscle gene expression (Mar and Ordahl, 1988, Gessler et al., 2024). In contrast, genes such as *Irx4*, *Gli1*, and *Dlx5* may be regulated indirectly through TEAD1-dependent transcriptional networks that control ventricular gene programs. Together, these findings indicate that *Tead1* mis-splicing likely compromises a TEAD1-dependent sarcomeric and ventricular gene network, thereby disrupting trabeculation and subsequent myocardial compaction.

Together, our findings support a model in which precise regulation of *Tead1* exon usage is required to maintain tissue-specific transcriptional outputs, and in which partial spliceosomal dysfunction caused by impaired PRPF8 activity selectively reduces TEAD1 function without complete protein loss. In the developing heart, this hypomorphic TEAD1 state disrupts processes underlying ventricular trabeculation and compaction. Importantly, although TEAD1 emerges as a major regulatory node linking spliceosomal dysfunction to ventricular wall morphogenesis, it is unlikely to be the sole downstream effector. Instead, partial spliceosome impairment appears to perturb a restricted set of key developmental regulators, thereby disrupting interconnected transcriptional, signalling, and morphogenetic programmes that collectively drive CHD.

## Acknowledgements

We thank Monica Justice for the l11Jus27 mouse line and Nkx2.5 *in situ* hybridization probe. We thank Benoit G Bruneau for the Irx4 *in situ* hybridization probe. We thank Ray Boot-Handfors for the Laminin antibody. Isabel foezoe for performing the genotyping.

We thank the University of Manchester Biological Services Facility for managing the mouse lines and the BioImaging Core Facility for the assistance with confocal microscopy.

## Author Contributions

WMSQ, KEH, HZ, AB, NA, AvdZ, MB, LAS, KJ, BB, DP performed research. WMSQ and KEH wrote original manuscript draft. KEH and CAJ provided supervision. KEH, CAJ and BB obtained funding. All authors read and edited the manuscript draft and approved submission.

## References

Ahmed, R. E., Tokuyama, T., Anzai, T., Chanthra, N. & Uosaki, H. 2022. Sarcomere maturation: function acquisition, molecular mechanism, and interplay with other organelles. Philosophical Transactions of the Royal Society B: Biological Sciences, 377.

Akerberg, B. N., Gu, F., Vandusen, N. J., Zhang, X., Dong, R., Li, K., Zhang, B., Zhou, B., Sethi, I., Ma, Q., Wasson, L., Wen, T., Liu, J., Dong, K., Conlon, F. L., Zhou, J., Yuan, G. C., Zhou, P. & Pu, W. T. 2019. A reference map of murine cardiac transcription factor chromatin occupancy identifies dynamic and conserved enhancers. Nat Commun, 10, 4907.

Atkinson, R., Georgiou, M., Yang, C., Szymanska, K., Lahat, A., Vasconcelos, E. J. R., Ji, Y., Moya Molina, M., Collin, J., Queen, R., Dorgau, B., Watson, A., Kurzawa-Akanbi, M., Laws, R., Saxena, A., Shyan Beh, C., Siachisumo, C., Goertler, F., Karwatka, M., Davey, T., Inglehearn, C. F., Mckibbin, M., Lührmann, R., Steel, D. H., Elliott, D. J., Armstrong, L., Urlaub, H., Ali, R. R., Grellscheid, S. N., Johnson, C. A., Mozaffari-Jovin, S. & Lako, M. 2024. PRPF8-mediated dysregulation of hBrr2 helicase disrupts human spliceosome kinetics and 5′-splice-site selection causing tissue-specific defects. Nat Commun, 15, 3138.

Bernier, Francois P., Caluseriu, O., Ng, S., Schwartzentruber, J., Buckingham, Kati J., Innes, A. M., Jabs, Ethylin W., Innis, Jeffrey W., Schuette, Jane L., Gorski, Jerome L., Byers, Peter H., Andelfinger, G., Siu, V., Lauzon, J., Fernandez, Bridget A., Mcmillin, M., Scott, Richard H., Racher, H., Majewski, J., Nickerson, Deborah A., Shendure, J., Bamshad, Michael J. & Parboosingh, Jillian S. 2012. Haploinsufficiency of *SF3B4*, a Component of the Pre-mRNA Spliceosomal Complex, Causes Nager Syndrome. The American Journal of Human Genetics, 90, 925-933.

Calses, P. C., Pham, V. C., Guarnaccia, A. D., Choi, M., Verschueren, E., Bakker, S. T., Pham, T. H., Hinkle, T., Liu, C., Chang, M. T., Kljavin, N., Bakalarski, C., Haley, B., Zou, J., Yan, C., Song, X., Lin, X., Rowntree, R., Ashworth, A., Dey, A. & Lill, J. R. 2023. TEAD Proteins Associate With DNA Repair Proteins to Facilitate Cellular Recovery From DNA Damage. Mol Cell Proteomics, 22, 100496.

Camenisch, T. D., Spicer, A. P., Brehm-Gibson, T., Biesterfeldt, J., Augustine, M. L., Calabro, A., Jr., Kubalak, S., Klewer, S. E. & Mcdonald, J. A. 2000. Disruption of hyaluronan synthase-2 abrogates normal cardiac morphogenesis and hyaluronan-mediated transformation of epithelium to mesenchyme. J Clin Invest, 106, 349–60.

Chen, L., Chan, S. W., Zhang, X., Walsh, M., Lim, C. J., Hong, W. & Song, H. 2010. Structural basis of YAP recognition by TEAD4 in the hippo pathway. Genes Dev, 24, 290–300.

Chen, Z., Friedrich, G. A. & Soriano, P. 1994. Transcriptional enhancer factor 1 disruption by a retroviral gene trap leads to heart defects and embryonic lethality in mice. Genes Dev, 8, 2293–301.

Chhatwal, K., Smith, J. J., Bola, H., Zahid, A., Venkatakrishnan, A. & Brand, T. 2023. Uncovering the Genetic Basis of Congenital Heart Disease: Recent Advancements and Implications for Clinical Management. CJC Pediatr Congenit Heart Dis, 2, 464–480.

Chiang, I. K. N., Humphrey, D., Mills, R. J., Kaltzis, P., Pachauri, S., Graus, M., Saha, D., Wu, Z., Young, P., Sim, C. B., Davidson, T., Hernandez-Garcia, A., Shaw, C. A., Renwick, A., Scott, D. A., Porrello, E. R., Wong, E. S., Hudson, J. E., Red-Horse, K., Del Monte-Nieto, G. & Francois, M. 2023. Sox7-positive endothelial progenitors establish coronary arteries and govern ventricular compaction. The EMBO Reports, 24, EMBR202255043.

Cowan, J. R. & Ware, S. M. 2015. Genetics and genetic testing in congenital heart disease. Clin Perinatol, 42, 373–93, ix.

Dupont, S., Morsut, L., Aragona, M., Enzo, E., Giulitti, S., Cordenonsi, M., Zanconato, F., Le Digabel, J., Forcato, M., Bicciato, S., Elvassore, N. & Piccolo, S. 2011. Role of YAP/TAZ in mechanotransduction. Nature, 474, 179–183.

Engal, E., Sharma, A., Aviel, U., Taqatqa, N., Juster, S., Jaffe-Herman, S., Bentata, M., Geminder, O., Gershon, A., Lewis, R., Kay, G., Hecht, M., Epsztejn-Litman, S., Gotkine, M., Mouly, V., Eiges, R., Salton, M. & Drier, Y. 2024. DNMT3B splicing dysregulation mediated by SMCHD1 loss contributes to DUX4 overexpression and FSHD pathogenesis. Science Advances, 10, eadn7732.

Fu, M., Hu, Y., Lan, T., Guan, K.-L., Luo, T. & Luo, M. 2022. The Hippo signalling pathway and its implications in human health and diseases. Signal Transduction and Targeted Therapy, 7, 376.

Galej, W. P., Oubridge, C., Newman, A. J. & Nagai, K. 2013. Crystal structure of Prp8 reveals active site cavity of the spliceosome. Nature, 493, 638–43.

García-Moreno, J. F. & Romão, L. 2020. Perspective in Alternative Splicing Coupled to Nonsense-Mediated mRNA Decay. Int J Mol Sci, 21.

Gessler, L., Siudzińska, A., Prószyński, T. J., Sandri, M., Von Eyss, B. & Hashemolhosseini, S. 2024. Deciphering the regulatory pathways in skeletal muscle lineage organized by the YAP1/TAZ-TEAD transcriptional network. bioRxiv, 2024.06.11.598443.

Goos, J. A. C., Swagemakers, S. M. A., Twigg, S. R. F., Van Dooren, M. F., Hoogeboom, A. J. M., Beetz, C., Günther, S., Magielsen, F. J., Ockeloen, C. W., M, A. R.-A., Pfundt, R., Yntema, H. G., Van Der Spek, P. J., Stanier, P., Wieczorek, D., Wilkie, A. O. M., Van Den Ouweland, A. M. W., Mathijssen, I. M. J. & Hurst, J. A. 2017. Identification of causative variants in TXNL4A in Burn-McKeown syndrome and isolated choanal atresia. Eur J Hum Genet, 25, 1126–1133.

Gottardi, C. J. & Gumbiner, B. M. 2001. Adhesion signaling: how beta-catenin interacts with its partners. Curr Biol, 11, R792–4.

Grainger, R. J. & Beggs, J. D. 2005. Prp8 protein: at the heart of the spliceosome. Rna, 11, 533–57.

Graziotto, J. J., Farkas, M. H., Bujakowska, K., Deramaudt, B. M., Zhang, Q., Nandrot, E. F., Inglehearn, C. F., Bhattacharya, S. S. & Pierce, E. A. 2011. Three gene-targeted mouse models of RNA splicing factor RP show late-onset RPE and retinal degeneration. Invest Ophthalmol Vis Sci, 52, 190–8.

Grego-Bessa, J., Luna-Zurita, L., Del Monte, G., Bolós, V., Melgar, P., Arandilla, A., Garratt, A. N., Zang, H., Mukouyama, Y. S., Chen, H., Shou, W., Ballestar, E., Esteller, M., Rojas, A., Pérez-Pomares, J. M. & De La Pompa, J. L. 2007. Notch signaling is essential for ventricular chamber development. Dev Cell, 12, 415–29.

Guo, W., Schafer, S., Greaser, M. L., Radke, M. H., Liss, M., Govindarajan, T., Maatz, H., Schulz, H., Li, S., Parrish, A. M., Dauksaite, V., Vakeel, P., Klaassen, S., Gerull, B., Thierfelder, L., Regitz-Zagrosek, V., Hacker, T. A., Saupe, K. W., Dec, G. W., Ellinor, P. T., Macrae, C. A., Spallek, B., Fischer, R., Perrot, A., Özcelik, C., Saar, K., Hubner, N. & Gotthardt, M. 2012. RBM20, a gene for hereditary cardiomyopathy, regulates titin splicing. Nat Med, 18, 766–73.

Henrique, D., Adam, J., Myat, A., Chitnis, A., Lewis, J. & Ish-Horowicz, D. 1995. Expression of a Delta homologue in prospective neurons in the chick. Nature, 375, 787–90.

Holden, J. K. & Cunningham, C. N. 2018a. Targeting the Hippo Pathway and Cancer through the TEAD Family of Transcription Factors. Cancers (Basel), 10.

Holden, J. K. & Cunningham, C. N. 2018b. Targeting the Hippo Pathway and Cancer through the TEAD Family of Transcription Factors. Cancers, 10, 81.

Isaac, R., Bandyopadhyay, G., Rohm, T. V., Kang, S., Wang, J., Pokhrel, N., Sakane, S., Zapata, R., Libster, A. M., Vinik, Y., Berhan, A., Kisseleva, T., Borok, Z., Zick, Y., Telese, F., Webster, N. J. G. & Olefsky, J. M. 2024. TM7SF3 controls TEAD1 splicing to prevent MASH-induced liver fibrosis. Cell Metab, 36, 1030–1043.e7.

Israeli-Rosenberg, S., Manso, A. M., Okada, H. & Ross, R. S. 2014. Integrins and Integrin-Associated Proteins in the Cardiac Myocyte. Circulation Research, 114, 572–586.

Jang, M. Y., Patel, P. N., Pereira, A. C., Willcox, J. A. L., Haghighi, A., Tai, A. C., Ito, K., Morton, S. U., Gorham, J. M., Mckean, D. M., Depalma, S. R., Bernstein, D., Brueckner, M., Chung, W. K., Giardini, A., Goldmuntz, E., Kaltman, J. R., Kim, R., Newburger, J. W., Shen, Y., Srivastava, D., Tristani-Firouzi, M., Gelb, B. D., Porter, G. A., Seidman, C. E. & Seidman, J. G. 2023. Contribution of Previously Unrecognized RNA Splice-Altering Variants to Congenital Heart Disease. Circulation: Genomic and Precision Medicine, 16, 224–231.

Jiang, F., Boylan, M., Maxwell, D. W., Shaikh Qureshi, W. M., Rowlands, C. F., Tenin, G., Mitchell, K., Stephen, L. A., Vasconcelos, E. J. R., Wang, D., Chen, T., Zha, J., Liu, J., Althali, N., Leordean, D. V., Gallagher, M. T., Basu, B., Szymanska, K., Veeraghanta, A., Keavney, B., Humphries, M. J., Ellingford, J., Smith, D., Johnson, C. A., O’keefe, R. T., Roy, S. & Hentges, K. E. 2025. The RNA splicing factor PRPF8 is required for left-right organiser cilia function and determination of cardiac left-right asymmetry via regulation of *Arl13b* splicing. bioRxiv, 2025.05.22.654869.

Jiang, S. W., Desai, D., Khan, S. & Eberhardt, N. L. 2000. Cooperative binding of TEF-1 to repeated GGAATG-related consensus elements with restricted spatial separation and orientation. DNA Cell Biol, 19, 507–14.

Joshi, S., Davidson, G., Le Gras, S., Watanabe, S., Braun, T., Mengus, G. & Davidson, I. 2017. TEAD transcription factors are required for normal primary myoblast differentiation in vitro and muscle regeneration in vivo. PLoS Genet, 13, e1006600.

Keightley, M. C., Crowhurst, M. O., Layton, J. E., Beilharz, T., Markmiller, S., Varma, S., Hogan, B. M., De Jong-Curtain, T. A., Heath, J. K. & Lieschke, G. J. 2013. In vivo mutation of pre-mRNA processing factor 8 (Prpf8) affects transcript splicing, cell survival and myeloid differentiation. FEBS Lett, 587, 2150–7.

Kile, B. T., Hentges, K. E., Clark, A. T., Nakamura, H., Salinger, A. P., Liu, B., Box, N., Stockton, D. W., Johnson, R. L., Behringer, R. R., Bradley, A. & Justice, M. J. 2003. Functional genetic analysis of mouse chromosome 11. Nature, 425, 81–6.

Li, F., Negi, V., Yang, P., Lee, J., Ma, K., Moulik, M. & Yechoor, V. K. 2022. TEAD1 regulates cell proliferation through a pocket-independent transcription repression mechanism. Nucleic Acids Res, 50, 12723–12738.

Li, J., Miao, L., Shieh, D., Spiotto, E., Li, J., Zhou, B., Paul, A., Schwartz, R. J., Firulli, A. B., Singer, H. A., Huang, G. & Wu, M. 2016. Single-Cell Lineage Tracing Reveals that Oriented Cell Division Contributes to Trabecular Morphogenesis and Regional Specification. Cell Rep, 15, 158–170.

Lines, M. A., Huang, L., Schwartzentruber, J., Douglas, S. L., Lynch, D. C., Beaulieu, C., Guion-Almeida, M. L., Zechi-Ceide, R. M., Gener, B., Gillessen-Kaesbach, G., Nava, C., Baujat, G., Horn, D., Kini, U., Caliebe, A., Alanay, Y., Utine, G. E., Lev, D., Kohlhase, J., Grix, A. W., Lohmann, D. R., Hehr, U., Böhm, D., Majewski, J., Bulman, D. E., Wieczorek, D. & Boycott, K. M. 2012. Haploinsufficiency of a spliceosomal GTPase encoded by EFTUD2 causes mandibulofacial dysostosis with microcephaly. Am J Hum Genet, 90, 369–77.

Liu, R., Jagannathan, R., Li, F., Lee, J., Balasubramanyam, N., Kim, B. S., Yang, P., Yechoor, V. K. & Moulik, M. 2019. Tead1 is required for perinatal cardiomyocyte proliferation. PLoS One, 14, e0212017.

Liu, R., Jagannathan, R., Sun, L., Li, F., Yang, P., Lee, J., Negi, V., Perez-Garcia, E. M., Shiva, S., Yechoor, V. K. & Moulik, M. 2020. Tead1 is essential for mitochondrial function in cardiomyocytes. Am J Physiol Heart Circ Physiol, 319, H89–h99.

Liu, R., Lee, J., Kim, B. S., Wang, Q., Buxton, S. K., Balasubramanyam, N., Kim, J. J., Dong, J., Zhang, A., Li, S., Gupte, A. A., Hamilton, D. J., Martin, J. F., Rodney, G. G., Coarfa, C., Wehrens, X. H., Yechoor, V. K. & Moulik, M. 2017. Tead1 is required for maintaining adult cardiomyocyte function, and its loss results in lethal dilated cardiomyopathy. JCI Insight, 2.

Lynch, D. C., Revil, T., Schwartzentruber, J., Bhoj, E. J., Innes, A. M., Lamont, R. E., Lemire, E. G., Chodirker, B. N., Taylor, J. P., Zackai, E. H., Mcleod, D. R., Kirk, E. P., Hoover-Fong, J., Fleming, L., Savarirayan, R., Boycott, K., Mackenzie, A., Brudno, M., Bulman, D., Dyment, D., Majewski, J., Jerome-Majewska, L. A., Parboosingh, J. S., Bernier, F. P. & Care4rare, C. 2014. Disrupted auto-regulation of the spliceosomal gene SNRPB causes cerebro–costo–mandibular syndrome. Nature Communications, 5, 4483.

Ma, Q., Zhang, Y. H., Guo, W., Feng, K., Huang, T. & Cai, Y. D. 2024. Machine Learning in Identifying Marker Genes for Congenital Heart Diseases of Different Cardiac Cell Types. Life (Basel), 14.

Magico, A. C. & Bell, J. B. 2011. Identification of a classical bipartite nuclear localization signal in the Drosophila TEA/ATTS protein scalloped. PLoS One, 6, e21431.

Malinová, A., Cvačková, Z., Matějů, D., Hořejší, Z., Abéza, C., Vandermoere, F., Bertrand, E., Staněk, D. & Verheggen, C. 2017. Assembly of the U5 snRNP component PRPF8 is controlled by the HSP90/R2TP chaperones. J Cell Biol, 216, 1579–1596.

Manuel, J. M., Guilloy, N., Khatir, I., Roucou, X. & Laurent, B. 2023. Re-evaluating the impact of alternative RNA splicing on proteomic diversity. Frontiers in Genetics, 14.

Mar, J. H. & Ordahl, C. P. 1988. A conserved CATTCCT motif is required for skeletal muscle-specific activity of the cardiac troponin T gene promoter. Proc Natl Acad Sci U S A, 85, 6404–8.

Mehta, Z. & Touma, M. 2023. Post-Transcriptional Modification by Alternative Splicing and Pathogenic Splicing Variants in Cardiovascular Development and Congenital Heart Defects. Int J Mol Sci, 24.

Meyer, H. V., Dawes, T. J. W., Serrani, M., Bai, W., Tokarczuk, P., Cai, J., De Marvao, A., Henry, A., Lumbers, R. T., Gierten, J., Thumberger, T., Wittbrodt, J., Ware, J. S., Rueckert, D., Matthews, P. M., Prasad, S. K., Costantino, M. L., Cook, S. A., Birney, E. & O’regan, D. P. 2020. Genetic and functional insights into the fractal structure of the heart. Nature, 584, 589–594.

Mia, M. M. & Singh, M. K. 2019. The Hippo Signaling Pathway in Cardiac Development and Diseases. Front Cell Dev Biol, 7, 211.

Nguyen, T. H. D., Galej, W. P., Bai, X.-C., Savva, C. G., Newman, A. J., Scheres, S. H. W. & Nagai, K. 2015. The architecture of the spliceosomal U4/U6.U5 tri-snRNP. Nature, 523, 47-52.

O’grady, L., Schrier Vergano, S. A., Hoffman, T. L., Sarco, D., Cherny, S., Bryant, E., Schultz-Rogers, L., Chung, W. K., Sacharow, S., Immken, L. L., Holder, S., Blackwell, R. R., Buchanan, C., Yusupov, R., Lecoquierre, F., Guerrot, A.-M., Rodan, L., De Vries, B. B. A., Kamsteeg, E. J., Santos Simarro, F., Palomares-Bralo, M., Brown, N., Pais, L., Ferrer, A., Klee, E. W., Babovic-Vuksanovic, D., Rhodes, L., Person, R., Begtrup, A., Keller-Ramey, J., Santiago-Sim, T., Schnur, R. E., Sweetser, D. A. & Gold, N. B. 2022. Heterozygous variants in PRPF8 are associated with neurodevelopmental disorders. American Journal of Medical Genetics Part A, 188, 2750–2759.

Pagliari, S., Vinarsky, V., Martino, F., Perestrelo, A. R., Oliver De La Cruz, J., Caluori, G., Vrbsky, J., Mozetic, P., Pompeiano, A., Zancla, A., Ranjani, S. G., Skladal, P., Kytyr, D., Zdráhal, Z., Grassi, G., Sampaolesi, M., Rainer, A. & Forte, G. 2021. YAP–TEAD1 control of cytoskeleton dynamics and intracellular tension guides human pluripotent stem cell mesoderm specification. Cell Death & Differentiation, 28, 1193–1207.

Pobbati, Ajaybabu V., Chan, Siew W., Lee, I., Song, H. & Hong, W. 2012. Structural and Functional Similarity between the Vgll1-TEAD and the YAP-TEAD Complexes. Structure, 20, 1135–1140.

Richards, A. A. & Garg, V. 2010. Genetics of congenital heart disease. Curr Cardiol Rev, 6, 91–7.

Ridge, L. A., Mitchell, K., Al-Anbaki, A., Shaikh Qureshi, W. M., Stephen, L. A., Tenin, G., Lu, Y., Lupu, I. E., Clowes, C., Robertson, A., Barnes, E., Wright, J. A., Keavney, B., Ehler, E., Lovell, S. C., Kadler, K. E. & Hentges, K. E. 2017. Non-muscle myosin IIB (Myh10) is required for epicardial function and coronary vessel formation during mammalian development. PLoS Genet, 13, e1007068.

Růžičková, Š. & Staněk, D. 2017. Mutations in spliceosomal proteins and retina degeneration. RNA Biol, 14, 544–552.

Sandireddy, R., Cibi, D. M., Gupta, P., Singh, A., Tee, N., Uemura, A., Epstein, J. A. & Singh, M. K. 2019. Semaphorin 3E/PlexinD1 signaling is required for cardiac ventricular compaction. JCI Insight, 4.

Sedmera, D., Pexieder, T., Vuillemin, M., Thompson, R. P. & Anderson, R. H. 2000. Developmental patterning of the myocardium. The Anatomical Record, 258, 319–337.

Silva, A. C., Pereira, C., Fonseca, A. C. R. G., Pinto-Do-Ó, P. & Nascimento, D. S. 2021. Bearing My Heart: The Role of Extracellular Matrix on Cardiac Development, Homeostasis, and Injury Response. Frontiers in Cell and Developmental Biology, Volume 8–2020.

Song, S., Zhang, X., Huang, Z., Zhao, Y., Lu, S., Zeng, L., Cai, F., Wang, T., Pei, Z., Weng, X., Luo, W., Lu, H., Wei, Z., Wu, J., Yu, P., Shen, L., Zhang, X., Sun, A. & Ge, J. 2024. TEA domain transcription factor 1 (TEAD1) induces cardiac fibroblasts cells remodeling through BRD4/Wnt4 pathway. Signal Transduction and Targeted Therapy, 9, 45.

Spielmann, N., Miller, G., Oprea, T. I., Hsu, C.-W., Fobo, G., Frishman, G., Montrone, C., Haseli Mashhadi, H., Mason, J., Munoz Fuentes, V., Leuchtenberger, S., Ruepp, A., Wagner, M., Westphal, D. S., Wolf, C., Görlach, A., Sanz-Moreno, A., Cho, Y.-L., Teperino, R., Brandmaier, S., Sharma, S., Galter, I. R., Östereicher, M. A., Zapf, L., Mayer-Kuckuk, P., Rozman, J., Teboul, L., Bunton-Stasyshyn, R. K. A., Cater, H., Stewart, M., Christou, S., Westerberg, H., Willett, A. M., Wotton, J. M., Roper, W. B., Christiansen, A. E., Ward, C. S., Heaney, J. D., Reynolds, C. L., Prochazka, J., Bower, L., Clary, D., Selloum, M., Bou About, G., Wendling, O., Jacobs, H., Leblanc, S., Meziane, H., Sorg, T., Audain, E., Gilly, A., Rayner, N. W., Aguilar-Pimentel, J. A., Becker, L., Garrett, L., Hölter, S. M., Amarie, O. V., Calzada-Wack, J., Klein-Rodewald, T., Da Silva-Buttkus, P., Lengger, C., Stoeger, C., Gerlini, R., Rathkolb, B., Mayr, D., Seavitt, J., Gaspero, A., Green, J. R., Garza, A., Bohat, R., Wong, L., Mcelwee, M. L., Kalaga, S., Rasmussen, T. L., Lorenzo, I., Lanza, D. G., Samaco, R. C., Veeraragaven, S., Gallegos, J. J., Kašpárek, P., Petrezsélyová, S., King, R., Johnson, S., Cleak, J., Szkoe-Kovacs, Z., Codner, G., Mackenzie, M., Caulder, A., Kenyon, J., Gardiner, W., Phelps, H., Hancock, R., Norris, C., Moore, M. A., Seluke, A. M., Urban, R., Kane, C., Goodwin, L. O., Peterson, K. A., Mckay, M., et al. 2022. Extensive identification of genes involved in congenital and structural heart disorders and cardiomyopathy. Nature Cardiovascular Research, 1, 157–173.

Tschaharganeh, D. F., Chen, X., Latzko, P., Malz, M., Gaida, M. M., Felix, K., Ladu, S., Singer, S., Pinna, F., Gretz, N., Sticht, C., Tomasi, M. L., Delogu, S., Evert, M., Fan, B., Ribback, S., Jiang, L., Brozzetti, S., Bergmann, F., Dombrowski, F., Schirmacher, P., Calvisi, D. F. & Breuhahn, K. 2013. Yes-Associated Protein Up-regulates Jagged-1 and Activates the NOTCH Pathway in Human Hepatocellular Carcinoma. Gastroenterology, 144, 1530–1542.e12.

Van Der Linde, D., Konings, E. E., Slager, M. A., Witsenburg, M., Helbing, W. A., Takkenberg, J. J. & Roos-Hesselink, J. W. 2011. Birth prevalence of congenital heart disease worldwide: a systematic review and meta-analysis. J Am Coll Cardiol, 58, 2241–7.

Verderame, M., Alcorta, D., Egnor, M., Smith, K. & Pollack, R. 1980. Cytoskeletal F-actin patterns quantitated with fluorescein isothiocyanate-phalloidin in normal and transformed cells. Proc Natl Acad Sci U S A, 77, 6624–8.

Verma, S. K., Deshmukh, V., Thatcher, K., Belanger, K. K., Rhyner, A. M., Meng, S., Holcomb, R. J., Bressan, M., Martin, J. F., Cooke, J. P., Wythe, J. D., Widen, S. G., Lincoln, J. & Kuyumcu-Martinez, M. N. 2022. RBFOX2 is required for establishing RNA regulatory networks essential for heart development. Nucleic Acids Res, 50, 2270–2286.

Wahl, M. C., Will, C. L. & Lührmann, R. 2009. The Spliceosome: Design Principles of a Dynamic RNP Machine. Cell, 136, 701–718.

Watkins, S. J., Borthwick, G. M. & Arthur, H. M. 2011. The H9C2 cell line and primary neonatal cardiomyocyte cells show similar hypertrophic responses in vitro. In Vitro Cellular & Developmental Biology - Animal, 47, 125–131.

Wen, T., Liu, J., He, X., Dong, K., Hu, G., Yu, L., Yin, Q., Osman, I., Peng, J., Zheng, Z., Xin, H., Fulton, D., Du, Q., Zhang, W. & Zhou, J. 2019. Transcription factor TEAD1 is essential for vascular development by promoting vascular smooth muscle differentiation. Cell Death Differ, 26, 2790–2806.

Wilkinson, E., Cui, Y.-H. & He, Y.-Y. 2022. Roles of RNA Modifications in Diverse Cellular Functions. Frontiers in Cell and Developmental Biology, 10.

Wood, K. A., Rowlands, C. F., Qureshi, W. M. S., Thomas, H. B., Buczek, W. A., Briggs, T. A., Hubbard, S. J., Hentges, K. E., Newman, W. G. & O’keefe, R. T. 2019. Disease modeling of core pre-mRNA splicing factor haploinsufficiency. Hum Mol Genet, 28, 3704–3723.

Wu, M. 2018. Mechanisms of Trabecular Formation and Specification During Cardiogenesis. Pediatr Cardiol, 39, 1082–1089.

Wu, W., He, J. & Shao, X. 2020. Incidence and mortality trend of congenital heart disease at the global, regional, and national level, 1990-2017. Medicine (Baltimore), 99, e20593.

Yamada, S., Ko, T., Ito, M., Sassa, T., Nomura, S., Okuma, H., Sato, M., Imasaki, T., Kikkawa, S., Zhang, B., Yamada, T., Seki, Y., Fujita, K., Katoh, M., Kubota, M., Hatsuse, S., Katagiri, M., Hayashi, H., Hamano, M., Takeda, N., Morita, H., Takada, S., Toyoda, M., Uchiyama, M., Ikeuchi, M., Toyooka, K., Umezawa, A., Yamanishi, Y., Nitta, R., Aburatani, H. & Komuro, I. 2023. TEAD1 trapping by the Q353R–Lamin A/C causes dilated cardiomyopathy. Science Advances, 9, eade7047.

Yang, M., Liu, Y., Lin, Z., Sun, H. & Hu, T. 2022. A novel de novo missense mutation in EFTUD2 identified by whole-exome sequencing in mandibulofacial dysostosis with microcephaly. J Clin Lab Anal, 36, e24440.

Yoshida, T. 2008a. MCAT elements and the TEF-1 family of transcription factors in muscle development and disease. Arterioscler Thromb Vasc Biol, 28, 8–17.

Yoshida, T. 2008b. MCAT Elements and the TEF-1 Family of Transcription Factors in Muscle Development and Disease. Arteriosclerosis, Thrombosis, and Vascular Biology, 28, 8–17.

Yue, W., Xiaosi, J., Yuhao, Z., Jing, Z. & Rulai, Y. 2021. Genetic and epigenetic mechanisms in the development of congenital heart diseases. World Journal of Pediatric Surgery, 4, e000196.

Zhang, W., Chen, H., Qu, X., Chang, C. P. & Shou, W. 2013. Molecular mechanism of ventricular trabeculation/compaction and the pathogenesis of the left ventricular noncompaction cardiomyopathy (LVNC). Am J Med Genet C Semin Med Genet, 163c, 144-56.

