## Supplementary files for "*Prpf8^N1531S^* homozygous mutant mouse embryos have multiple defects in cardiac development and show aberrant splicing of the cardiac transcription factor *Tead1*"

**Supplementary Table 1: List of antibodies**

| **Antibodies** | **Supplier and catalogue number** | **Dilution** |
| --- | --- | --- |
| MF20 | DHSB (MF20) | 1:100 |
| Endomucin | SantaCruz (Sc-65495) | 1:100 |
| PECAM | BD Biosciences (550274) | 1:100 |
| Prpf8 | Abcam (ab79237) | 1:200 |
| Anti-Flag | Simga Aldrich (F3165) | 1:200 |
| α-Actinin | Proteintech (11313-2-AP) | 1:200 |
| Arl13B | Proteintech (17711-1-AP) | 1:200 |
| Fibronectin 1 | Proteintech (15613-1-AP) | 1:200 |
| Laminin | Abcam ab11575 | 1:200 |
| Collagen IV (ColIV) | Abcam (ab19808) | 1:200 |
| Non-Muslce Myosin IIB (NMIIB) | BioLegend (PRB-445P) | 1:200 |
| β-Catenin | Proteintech (51067-2-AP) | 1:200 |
| Wheat Germ Agglutinin (WGA)-568 | Biotium (29077-1) | 1:500 |
| Phalloidin 488 | Thermo Fischer Scientific (A12379) | 1:400 |
| Phospho-Histone 3 | Proteintech (17168-1-AP) | 1:200 |
| Hyaluronan Synthase 2 (Has2) | Bioryt (Orb15430) | 1:200 |
| NICD1 | Cell signalling (7194) | 1:100 |
| β-Catenin | Proteintech (51067-2-AP) | 1:200 |
| Alexa Flour® 488 | Thermo Fischer Scientific (A-11001) | 1:500 |
| Alexa Flour® 568 | Thermo Fischer Scientific (A-11011) | 1:500 |
| Alexa Flour® 647 | Thermo Fischer Scientific (A-21247) | 1:500 |
| DAPI | Sigma Aldrich (D9542) | 1:500 |

**Supplementary table 2: List of RT-PCR primers sequence.**

| Primer Name | Primer sequence |
| --- | --- |
| Tead1 exon 2 F | TGAGCAGAGTTTCCAGGAGG |
| Tead1 exon 3 R | TCCTGGTCCTTGTCTTTCCC |
| Tead1 exon 7 R | GCTTGTTGTGGATGGCAGTA |
| Tead1 exon 10 R | AGGCTCAAACCCTGGAATGG |

**Supplementary table 3: List of in situ hybridisation probe primer sequences**

The red highlighted sequence represents either the T3 or T7 primer sequence.

| Probe | Sequence |
| --- | --- |
| Tnnt1 T3 | AAATTAACCCTCACTAAAGGGGAAGCGCATGGAGAAAGAC |
| Tnnt1 T7 | TAATACGACTCACTATAGGGTGTCCTGGCAGTCTCACTTC |
| Myl2 T3 | AAATTAACCCTCACTAAAGGCAAGAAGCGGATAGAAGGCG |
| Myl2 T7 | TAATACGACTCACTATAGGGGACCACCATCAAGCCTTGTG |
| Myl4 | AAATTAACCCTCACTAAAGGGCAACCGACAGTGTCCATATA |
| Myl4 | TAATACGACTCACTATAGGGGAACTCATCTCTTCAGGCTTG |
| Gli1 T3 | AAATTAACCCTCACTAAAGGCCTTCTCATGCTGGGGTGTA |
| Gli1 T7 | TAATACGACTCACTATAGGGCAGAGGGAGATGGGGTGTTT |
| Dlx5 T3 | AAATTAACCCTCACTAAAGGCCAGAGGTGAGGATGGTGAA |
| Dlx5 T7 | TAATACGACTCACTATAGGGGATAGTGTCCACAGTTGCGC |
